# Using adversarial networks to extend brain computer interface decoding accuracy over time

**DOI:** 10.1101/2022.08.26.504777

**Authors:** Xuan Ma, Fabio Rizzoglio, Eric J. Perreault, Lee E. Miller, Ann Kennedy

## Abstract

Existing intracortical brain computer interfaces (iBCIs) transform neural activity into control signals capable of restoring movement to persons with paralysis. However, the accuracy of the “decoder” at the heart of the iBCI typically degrades over time due to turnover of recorded neurons. To compensate, decoders can be recalibrated, but this requires the user to spend extra time and effort to provide the necessary data, then learn the new dynamics. As the recorded neurons change, one can think of the underlying movement intent signal being expressed in changing coordinates. If a mapping can be computed between the different coordinate systems, it may be possible to stabilize the original decoder’s mapping from brain to behavior without recalibration. We previously proposed a method based on Generalized Adversarial Networks (GANs), called “Adversarial Domain Adaptation Network” (ADAN), which aligns the distributions of latent signals within underlying low-dimensional neural manifolds. However, ADAN was tested on only a very limited dataset. Here we propose a method based on Cycle-Consistent Adversarial Networks (Cycle-GAN), which aligns the distributions of the full-dimensional neural recordings. We tested both Cycle-GAN and ADAN on data from multiple monkeys and behaviors and compared them to a linear method based on Procrustes Alignment of axes provided by Factor Analysis (PAF). Both GAN-based methods outperformed PAF. Cycle-GAN and ADAN (like PAF) are unsupervised and require little data, making them practical in real life. Overall, Cycle-GAN had the best performance and was easier to train and more robust than ADAN, making it ideal for stabilizing iBCI systems over time.

**Significance Statement:** The inherent instabilities in the neural signals acquired by intracortical microelectrode arrays cause the performance of an intracortical brain computer interface (iBCI) decoder to drop over time, as the movement intent signal must essentially be recorded from neurons representing an ever-changing coordinate system. Here, we address this problem using Generative Adversarial Networks (GANs) to align these coordinates and compare their success to another, recently proposed linear method that uses Factor Analysis and Procrustes alignment. Our proposed methods are fully unsupervised, can be trained quickly, and require remarkably little new data. These methods should give iBCI users access to decoders with unchanging dynamics, and without the need for periodic supervised recalibration.

## Introduction

Intracortical brain-computer interfaces (iBCIs) aim to restore motor function in people with paralysis by transforming neural activity recorded from motor areas of the brain into an estimate of the user’s movement intent. This transformation is accomplished using a neural “decoder”, an algorithm that translates the moment-to-moment activity of a population of neurons into a signal used to control intended movements. There has been substantial improvement in our ability to record and decode from large populations of neurons in the past decade, which allows more information to be extracted from the brain and conveyed to the external effectors of the iBCI. However, the long-term stability of iBCIs is still far from satisfactory due in part to the instabilities in neural recordings. The relative micromotion between the electrode tip and the brain tissue (1), the changes of regional extracellular environment (2), or even the active and inactive state shifts of neurons (3) could contribute to such instabilities, resulting in the turnover of signals picked by the chronically implanted electrodes on a time scale of days or even a few hours (4). Given these changes, a decoder could produce inaccurate predictions of the user’s intent leading to the degraded iBCI performance.

To counteract these effects, a neural decoder might be recalibrated with newly acquired data. A disadvantage of this strategy is that during recalibration, normal use would be interrupted. Furthermore, the recalibration process likely means the user would need to learn the dynamics of the new decoder, imposing additional time and cognitive burden. For persons with paralysis to live more independently, an ideal iBCI would accommodate the gradual drift in neural recordings without supervision, thereby minimizing the need to periodically learn new decoders. For the performance of the initial “day-0” decoder to be maintained, an additional component, an “input stabilizer”, would need to be added to transform the neural recordings made on a later day (“day-k”) such that they take on the statistics of the day-0 recordings.

Recently there has been a great deal of interest in the concept of a low-dimensional neural manifold embedded within the neural space that is defined by the full set of recorded neurons, and the “latent signals” that can be computed in it (5). A previous paper from our group demonstrated that by aligning the day-k and day-0 latent signals using canonical correlation analysis (CCA), the performance of a fixed day-0 decoder could be maintained over months and even years, despite turnover of the neural recordings.

Unfortunately, CCA has a couple significant limitations. For one, it is a linear process, not able to account for the nonlinear mappings that have been demonstrated between high-dimensional neural recordings and their low-dimensional manifolds (6, 7). Also, its use in a real-life scenario would be cumbersome. This application of CCA can be thought of as rotating two sets of neural signals “spatially” to achieve optimal overlap (and temporal correlation). To do so requires that the behaviors on day-0 and day-k have the same temporal structure. If they do not, no amount of spatial rotation will achieve a correlation between the neural signals. However, motor behaviors in daily life are typically not well structured, with well-defined onsets and offsets, making time alignment difficult, if not impossible. Where this method has been used successfully, it has been with highly stereotypic behaviors with distinct trial structure.

Another recently published linear method for decoder stabilization uses a Procrustes-based (8) alignment on low-dimensional manifolds obtained from the neural activity using Factor Analysis (9). This approach, which we will refer to as “Procrustes Alignment of Factors” (PAF), successfully stabilized online iBCI cursor control with a fixed decoder. Time alignment is not needed for PAF, as it aligns the coordinate axes for the manifolds directly. However, it does require a subset of the coordinate axes in which the manifold is embedded (the neural recording channels) to be unchanged between days 0 and k. Furthermore, the use of a Procrustes-based transformation means that this strategy cannot correct for nonlinear changes in the neural manifold across days.

In another approach to decoder stabilization, we view changes in neural recordings as arbitrary shifts in the distribution of population firing rates. From this perspective, the reason for poor cross-day performance of decoders is clear: a decoder that is trained only on observations from a given distribution (e.g., those of “day-0”) won’t perform well on data from other distributions (i.e., “day-k”). A machine learning approach termed “domain adaptation” has been used to cope with such distribution mismatches by learning a transformation that minimizes the difference between the transformed distributions; this permits a model trained on one distribution (the source domain) to generalize to the other, the target domain (10, 11). For example, if we have a classifier trained to distinguish photos of objects, domain adaptation could be used to transform drawings of those objects into “photo-like” equivalents, so that our existing photo-based classifier can be used. Domain adaptation can be implemented with Generative Adversarial Networks (GANs; (12)). GANs use two networks – a generator trained to transform a source distribution into a target distribution, and a discriminator trained to do the opposite: determine whether a given distribution is real or synthesized by the generator. The adversarial nature of the generator and discriminator enables the model to be trained in an unsupervised manner (13, 14). GAN-based domain adaptation has been applied to computer vision problems, like adapting a classifier trained to recognize the digits of one style for use in recognizing those of another style (14), or translating images in the style of one domain to another (e.g., colorizing black-and-white photos (15)).

We recently developed an approach we named Adversarial Domain Adaptation Network (ADAN; (16)), that used a GAN to perform domain adaptation to enable a fixed day-0 iBCI decoder to work accurately on input signals recorded on day-k. ADAN finds low-dimensional manifolds using a nonlinear autoencoder, and aligns the empirical distribution of the day-k recordings (the source domain) to those of day-0 (the target domain) by aligning the distributions of residuals (as in (17)) between neural firing rates and their nonlinear autoencoder reconstructions (that is, the portion of neurons’ activity not predicted from the manifold). Note that, compared to PAF, ADAN performs the alignment in the high-dimensional space of reconstructed firing rates, but requires the computation of a low-dimensional manifold to do so. In the earlier study we found that ADAN outperforms both CCA and an alignment process that minimized the KL divergence between the distributions of the day-k and day-0 latent spaces (Kullback-Leibler Divergence Minimization, KLDM). However, ADAN was only tested on data from a single monkey and a single task, for just two weeks. Our subsequent exploration into applying ADAN to other datasets suggests that, while it can work in other settings, its performance is quite sensitive to model hyperparameter settings. This is consistent with previous reports that GANs can be highly dependent on choice of architecture and a variety of hyperparameter settings (18). We therefore sought alternative GAN-based approaches that might offer more robust performance.

Recently, Zhu et al. developed a novel GAN architecture named Cycle-Consistent Adversarial Networks (Cycle-GAN; (19)) in the context of image domain adaptation. Cycle-GAN introduced a mechanism termed “cycle-consistency”, which helps to regularize model performance. Specifically, Cycle-GAN implements both forward and inverse mappings between a pair of domains: the forward mapping translates data in the source domain to the target domain, while the inverse mapping brings the translated data back to the source domain. This regularization mechanism forces the learned transformation between the source and the target distributions to be a bijection, thereby reducing the search space of possible transformations (19, 20). In addition to its promise of robustness, an advantage of Cycle-GAN is that it can be used to align the full-dimensional distributions of the day-0 and day-k recordings without the need to compute an underlying manifold. All other alignment methods that have been explored (CCA, PAF, KLDM and ADAN) work with lowdimensional latent signals. This leads to the advantage that the (small) information loss caused by dimensionality reduction can be avoided. Also, as most existing iBCI decoders are computed directly from the full-dimensional neural recordings, no extra transformation would be required.

In this study, we compare Cycle-GAN, ADAN and PAF using datasets from several monkeys, spanning a broad variety of motor behaviors, and spanning several months. We chose not to test CCA, as it requires time alignment of the data, and it (as well as KLDM) was outperformed by ADAN in our earlier study (16). We found that both GAN-based methods outperformed PAF. We also demonstrated that the addition of cycle-consistency improved the alignment and made training much less dependent on hyper-parameters.

## Results

### Performance of a well-calibrated iBCI decoder declines over time

We trained six monkeys to perform five tasks: power and key grasping, center-out target tracking using isometric wrist torque, and center-out and random-target reaching movements (SI Appendix, Fig. S1). After training, each monkey was implanted with a 96-channel microelectrode array in either the hand or arm area of M1. Four animals (monkeys J, S, G, P) were also implanted with intramuscular leads in forearm and hand muscles contralateral to the cortical implant; these were used to record electromyograms (EMGs). We recorded multi-unit activity on each M1 electrode together with motor output (EMGs and/or hand trajectories) for many sessions across multiple days. All recording sessions for a specific task and an individual monkey were taken together to form a dataset. We collected a total of seven datasets, and the recording sessions in each of them spanned from ~30 to ~100 days (See Methods; SI Appendix, Table S1).

As in previous studies (1, 21), we found substantial instability in the M1 neurons we recorded over time, even though the motor outputs and task performance were generally stable (See SI Appendix, supplementary results; Fig. S2, Fig. S3). We first asked how this instability affected the performance of an iBCI decoder. We fit a Wiener filter decoder with data recorded on a reference day (designated “day-0”; Fig. 1A). We then used this decoder to predict the motor outputs from M1 neural recordings on later days (“day-k”) and computed the coefficient of determination (R^2^) between the predictions and the actual data (see Methods). Fig. 2 shows example predictions from each task. In all cases, both EMG (top row) and kinematic (bottom row) decoders could reconstruct movement trajectories with high accuracy on held-out trials from the day of training (“day-0”). However, the calibrated day-0 decoders consistently failed to predict EMGs or hand trajectories accurately on day-k. The degradation of the performance across time occurred for all behavioral tasks and monkeys, and could be substantial even a few days after decoder training (SI Appendix, Fig. S4).

**Figure 1.**
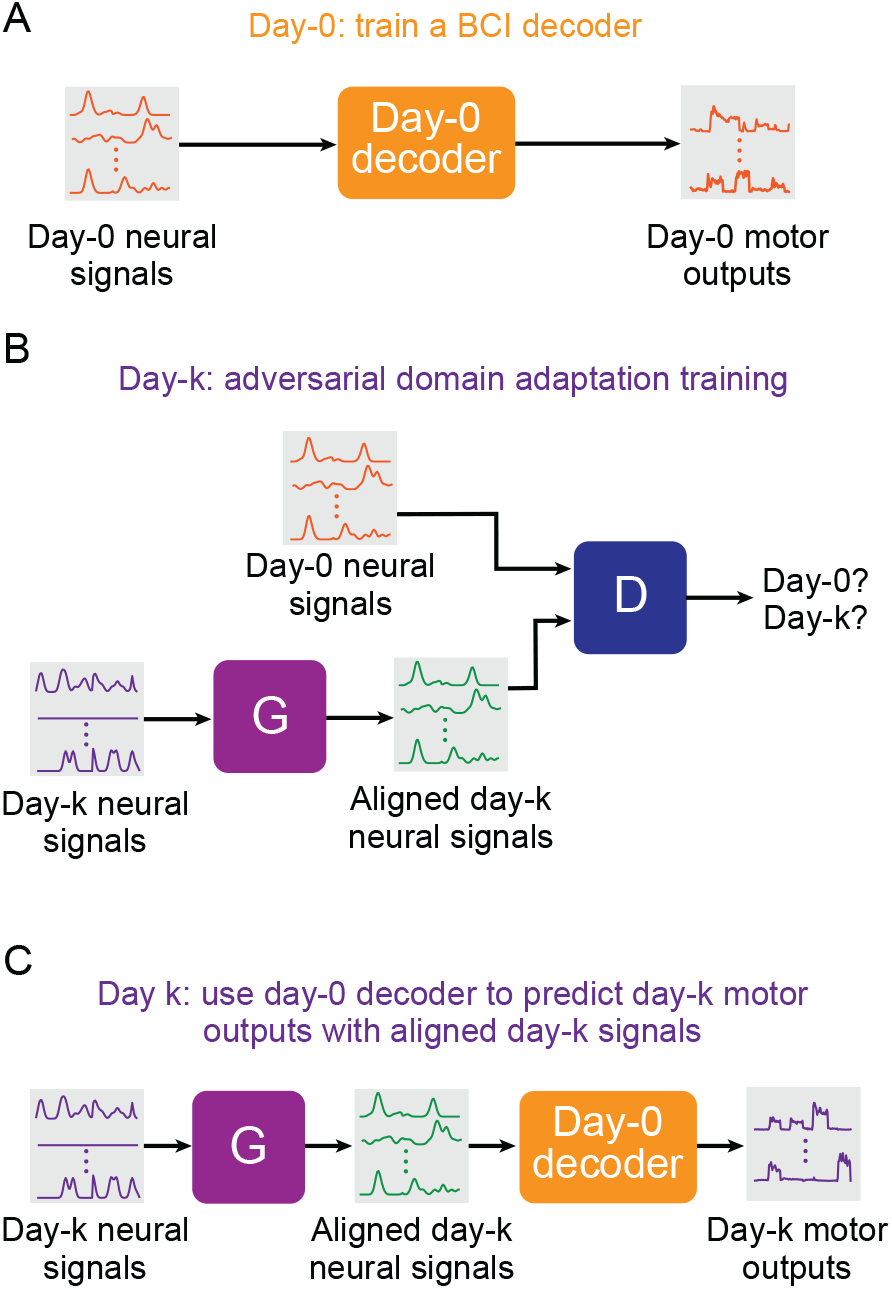
Setup for stabilizing an intracortical brain computer interface (iBCI) with adversarial domain adaptation. (*Λ*) Initial BCI decoder training on day-0. The decoder is computed to predict the motor outputs from neural signals, using either the full-dimensional neural recordings or the low-dimensional latent signals obtained through dimensionality reduction. This decoder will remain fixed over time after training. *(B)* A general framework for adversarial domain adaptation training on a subsequent day-k. The “Generator” (G) is a feedforward neural network that takes day-k neural signals as the inputs and aims to transform them into a form similar to day-0 signals; we also refer to G as the “aligner”. The “Discriminator” (D) is another feedforward neural network that takes both the outputs of G (aligned day-k neural signals) and day-0 neural signals as the inputs and aims to discriminate between them. (*C*) A trained aligner and the fixed day-0 decoder are used for iBCI decoding on day-k. The aligned signals generated by G are fed to the day-0 decoder to produce the predicted motor outputs.

**Figure 2.**
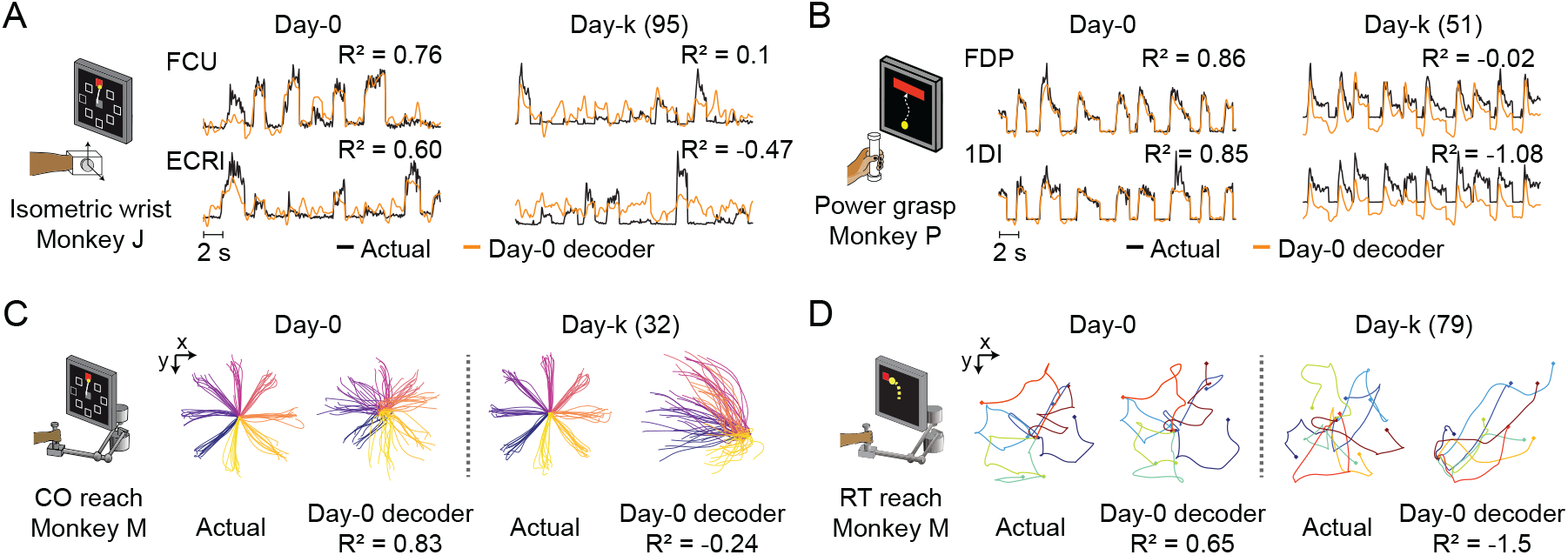
The performance of well-calibrated decoders declines over time. (*A*) Actual EMGs (black) and predicted EMGs (orange) using the day-0 decoder for flexor carpi ulnaris (FCU) and extensor carpi radialis longus (FCRl) during the isometric wrist task. (*B*) Actual and predicted EMGs using the day-0 decoder for flexor digitorum profundus (FDP) and first dorsal interosseous (1DI) during the power grasp task. (*C*) Actual hand trajectories and predictions using the day-0 decoder during the center-out (CO) reach task. Colors represent different reaching directions. *(D)* Actual and predicted hand trajectories using the day-0 decoder during the random-target (RT) reach task. Colors represent different reaching directions.

### Adversarial networks mitigate the performance declines of day-0 decoders

We proposed to use generative adversarial network (GAN) based domain adaptation (Fig. 1B) to address the problem described above. We tested two different architectures: Adversarial Domain Adaptation Network (ADAN) (16), and Cycle-Consistent Adversarial Networks (Cycle-GAN) (19). As both ADAN and Cycle-GAN were trained to reduce the discrepancy between the neural recordings on day-0 and those on day-k by aligning their probability density functions (PDFs), we call them “aligners”. We used the dataset with the longest recording timespan (monkey J, isometric wrist task, spanning 95 days) to determine appropriate choices of the hyperparameters for neural network training, which are presented in detail in a later section. We used the resulting hyperparameter values for the tests of all other monkeys and tasks. For comparison, we also used all datasets to test another type of “aligner” that aimed to align the low-dimensional neural manifolds between day-0 and day-k (9), which we termed “Procrustes Alignment of Factors” (PAF).

The tests were conducted with the procedures presented by Fig. 1. First, we picked a given day as day-0, and used the data recorded on that day to fit a Wiener filter as the “day-0 decoder” (Fig. 1A). Then, we trained the three types of aligners (ADAN, Cycle-GAN and PAF) to align the neural recordings on a different day (day-k) to those on day-0 (Fig. 1B). Each day in a dataset other than the designated day-0 was treated as a day-k, whether it occurred before or after day-0. Finally, we processed the neural recordings on day-k with the trained aligners, fed the aligned signals to the fixed day-0 decoder, and evaluated the accuracy of the predictions this decoder could obtain (Fig. 1C). For each of the seven datasets being tested, we repeated these three procedures for multiple instantiations using different day-0s (see SI Appendix, Table S1). To characterize the performance of the day-0 decoder after alignment, we represent the decoder accuracy as the “performance drop” with respect to a daily recalibrated decoder (R^2^_aligned_ – R^2^_same-day_). If an aligner works perfectly, we expect the performance drop of day-0 decoders to be close to 0, which means the decoder achieves accuracy equal to a within-day decoder after the alignment.

Unlike ADAN and PAF, Cycle-GAN alignment does not require computation of a latent representation from neural recordings. As a result, Cycle-GAN is naturally suited to a decoder trained on the full-dimensional neural firing rate signals. It is theoretically possible to use a full-dimensional decoder with ADAN and PAF as well, by training on firing rates reconstructed from the latent spaces of the ADAN autoencoder and PAF factors respectively. However, we found that the performance of these full-dimensional decoders was inferior to that of a decoder trained on the inferred latent signals (SI Appendix, Fig. S5). For completeness, we also tested a decoder trained on Cycle-GAN-generated firing rates projected into a low-dimensional manifold obtained using Factor Analysis; as expected, its performance was slightly worse than that of a full-dimensional decoder, but still better than ADAN and PAF with a low-dimensional decoder (SI Appendix, Fig. S5).

In light of the analysis above, we here compare the better-performing of the two potential decoder input formats for each alignment method: full-dimensional for Cycle-GAN, and low-dimensional for ADAN and PAF (see Methods for details). Aside from this difference of input dimensionality, the architecture of the day-0 decoder (a Wiener filter) was the same for all aligners. The accuracy of the day-0 decoders of the three aligners was modestly but significantly different across tasks: ADAN: R^2^ = 0.73 ± 0.009 (mean ± s.e.); Cycle-GAN: R^2^ = 0.72 ± 0.009; PAF: R^2^ = 0.71 ± 0.009 (P= 0.008, linear mixed-effect model with the type of aligner as fixed and the type of task as random factor).

We then fit a linear mixed-effect model with type of aligner and days as fixed factors and type of task as random factor for a quantitative evaluation of the performance of the three aligners. The accuracy of the day-0 decoder on data collected on the day immediately following day-0 (i.e., day-1) after alignment was significantly different across the aligners (Cycle-GAN: −0.02 ± 0.004 (mean ± s.e.); ADAN: −0.06 ± 0.005; PAF: −0.11 ± 0.005; P ~ 0). Cycle-GAN significantly outperformed both ADAN (P ~ 0) and PAF (P ~ 0). ADAN also significantly outperformed PAF (P ~ 0).

The performance degradation of day-0 decoders for periods greater than one day (SI Appendix, Fig. S4) was also mitigated by all three alignment methods, although to different extents. Nonetheless, there remained a significant and increasing performance drop over time (Fig. 3A and 3B). We found a significant interaction between time and alignment method (P = 0.026), indicating that there was a difference between methods in performance drop over time, and a post-hoc comparison showed that Cycle-GAN had the least overall performance degradation, significantly better than PAF, and better, but not significantly so, than ADAN (P = 0.008 vs PAF; P = 0.328 vs ADAN). ADAN was better, but not significantly, than PAF (P = 0.091). Taken together, this analysis shows that Cycle-GAN moderately outperforms both ADAN and PAF (see also Fig. 3C; SI Appendix, Fig. S6B and S6C), and furthermore that the two nonlinear alignment methods tend to be more stable over time than PAF (see also Fig. 3C; SI Appendix, Fig. S6A and S6B).

**Figure 3.**
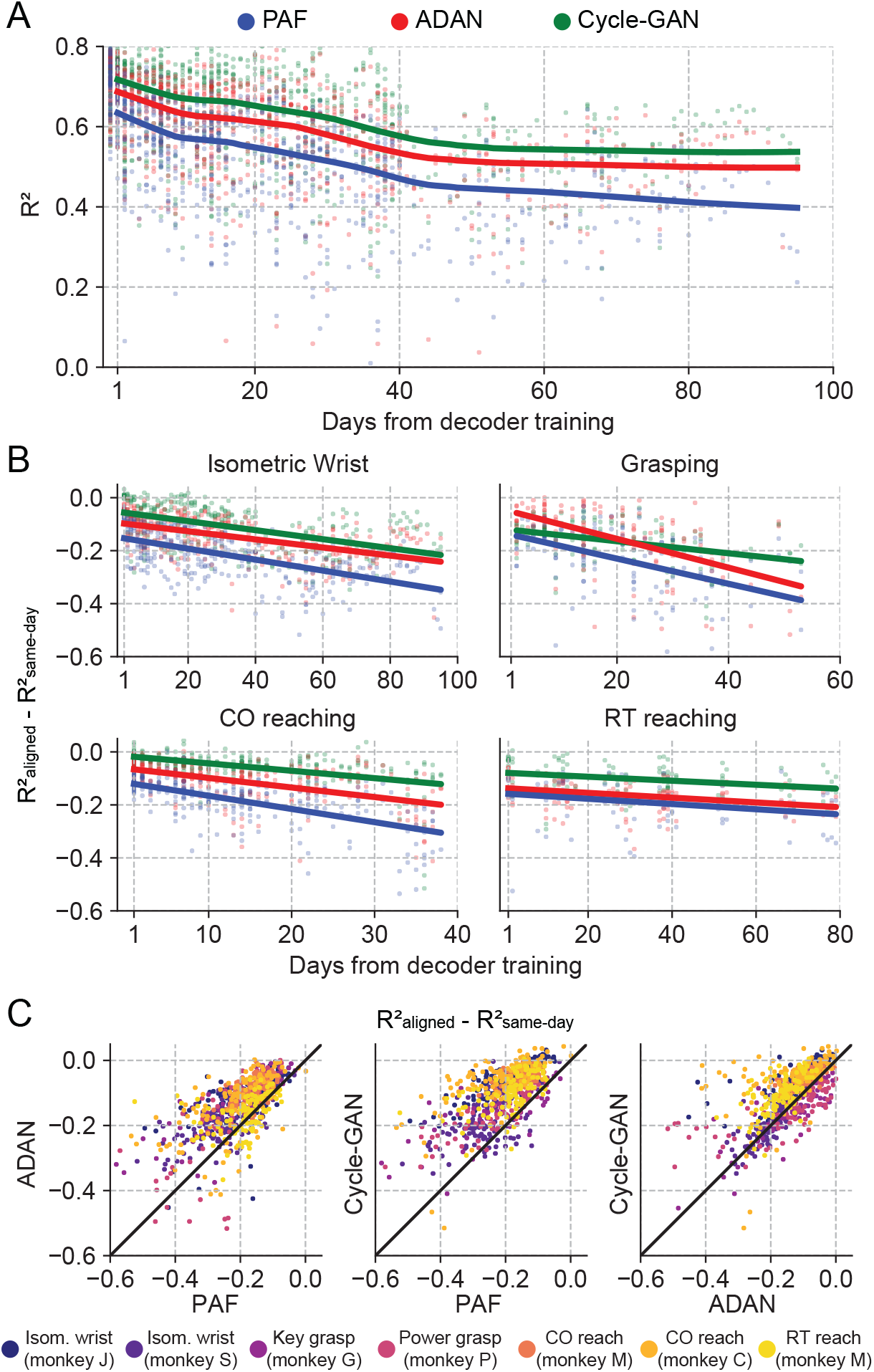
The proposed GANs-based domain adaptation methods outperform Procrustes Alignment of Factors in diverse experimental settings. (*A*) Prediction accuracy over time using the fixed decoder trained on day-0 data is shown for all experimental conditions (single dots: R^2^ as a function of days after decoder training, lines: locally weighted scatterplot smoothing fits). We compared the performance of the day-0 decoder after domain adaptation alignment with Cycle-GAN (green), ADAN (red) and PAF (blue). (*B*) We computed the prediction performance drop with respect to a daily-retrained decoder (single dots: R^2^ drop (R^2^_aligned_ - R^2^_same-day_) for days after decoder training, lines: linear fits). Cycle-GAN and ADAN both outperformed PAF, with Cycle-GAN degrading most slowly for all the experimental conditions. (*C*) We compared the performance of each pair of aligners by plotting the prediction performance drop of one aligner versus that of another. Each dot represents the R^2^ drop after decoder training relative to the within-day decoding. Marker colors indicate the task. Both proposed domain adaptation techniques outperformed PAF (left and center panels), with Cycle-GAN providing the best domain adaptation for most experimental conditions (right panel).

### Cycle-GAN is robust to hyperparameter settings

While they can be powerful, GANs can present a training challenge: choosing suitable hyperparameters is important, for example, to balance the learning process and prevent either of the two networks (the generator or discriminator) from dominating the loss function. High sensitivity of model performance to hyperparameter values would pose a potential barrier to the adoption of either ADAN or Cycle-GAN as a tool for cross-day alignment. As in (22), we assessed sensitivity to hyperparameters by testing the impact of batch size and learning rates on alignment performance. Because these hyperparameter sweeps are very computationally expensive, we evaluated them using only the single dataset with the greatest span of time.

We trained both ADAN and Cycle-GAN aligners on day-k data relative to four selected day-0 reference days. We kept the learning rates for the generator (LR_G_) and the discriminator (LR_D_) fixed (for ADAN, LR_G_ = 0.0001, LR_D_/LR_G_ = 0.5; for Cycle-GAN, LR_G_ = 0.0001, LR_D_/LR_G_ = 10). As in the previous section, we evaluated the drops in aligned day-0 decoder accuracy. We found that ADAN maintained good performance when batch size was small, but that performance started to drop significantly for larger batch sizes (64: −0.13 ± 0.0096 (mean ± s.e.); 256: −0.17 ± 0.013; P ~ 0, Wilcoxon’s signed rank test; Fig. 4A). In contrast, Cycle-GAN based aligners performed consistently at all tested batch sizes. These results suggest that ADAN may need a small batch size, while Cycle-GAN based aligners have no strong requirement.

**Figure 4.**
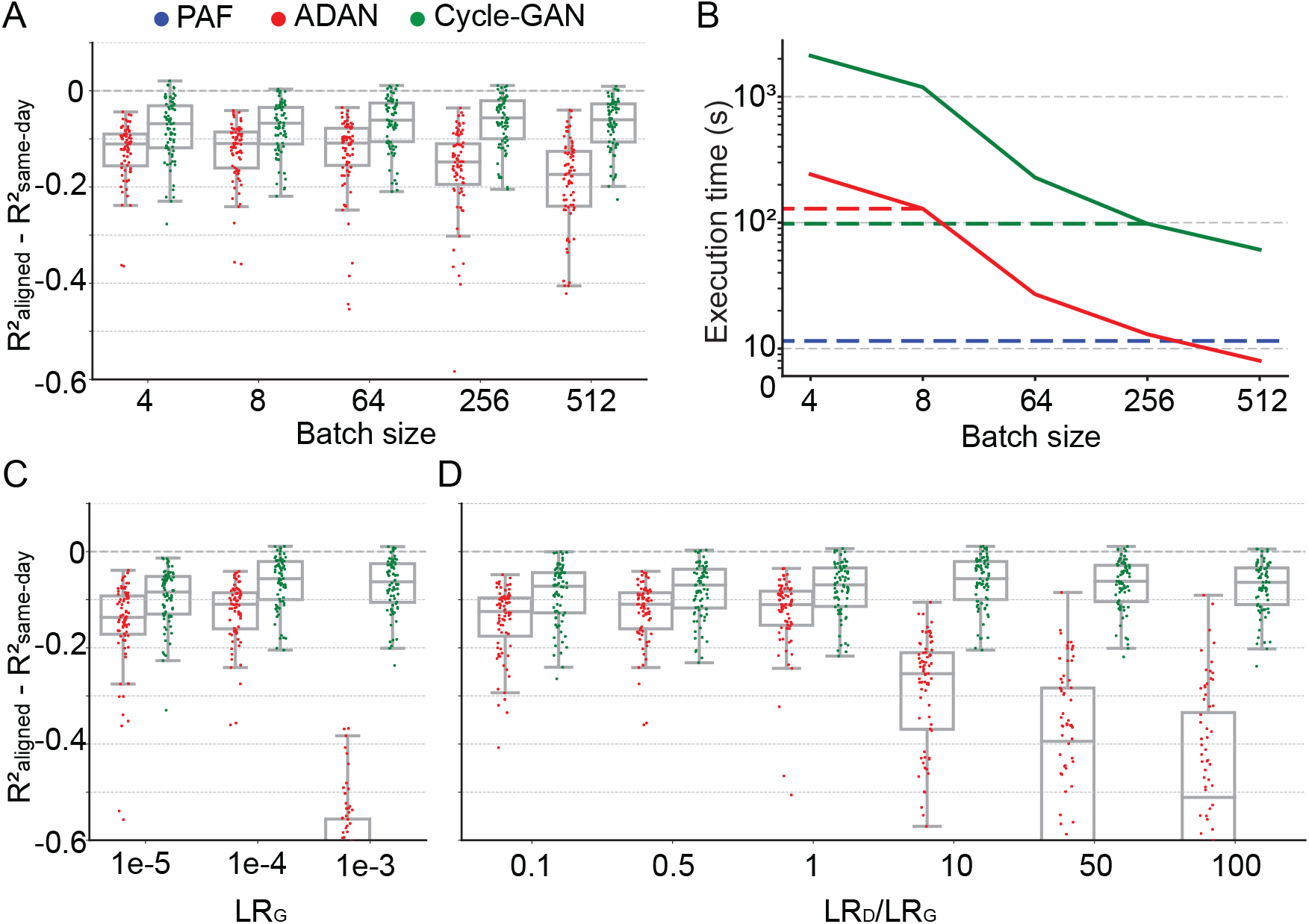
Cycle-GAN is more robust to hyperparameter tuning than ADAN. Effect of different batch sizes during training of Cycle-GAN (green) and ADAN (red) with mini-batch gradient descent on (*A*) the day-k performance of 4 selected day-0 decoders and (*B*) the execution time of 200 training epochs. The much faster execution time of PAF (blue) is also shown for reference. Compared to ADAN, Cycle-GAN did not require a small batch size, resulting in faster training (Cycle-GAN: 98s with batch size 256; ADAN: 129s with batch size 8; FA aligner: 11.5s). Effect of training each domain adaptation method with different generator (*C*) and discriminator (*D*) learning rate. The generator and the discriminator learning rate were denoted as LRG and LRD, respectively. For LRD testing, we kept LRG fixed (LRG = 1e-4 for both ADAN and Cycle-GAN), and changed the ratio between LRD and LRG (LRD/LRG). ADAN-based aligners did not perform well for large LRG or LRD/LRG values, while Cycle-GAN-based aligners remained stable for all the testing conditions. In (*A*), (*C*) and (*D*) single dots show the prediction performance drop on each day-k relative to the 4 selected day-0s with respect to the R^2^ of a daily-retrained decoder (R^2^_aligned_ - R^2^_same-day_). Boxplots show 25^th^, 50^th^ and 75^th^ percentiles of the R^2^ drop with the whiskers extending to the entire data spread, not including outliers.

Neural network training time is inversely proportional to batch size - therefore given two batch size options that give comparable model performance, the larger of the two will yield faster training. We found that Cycle-GAN was slower than ADAN for smaller batch sizes, although neither method required more than a few minutes when operating within their optimal batch size range (Fig. 4B). Thus, we set the ADAN batch size for subsequent analyses to 8 and for Cycle-GAN to 256. Although we could have increased the batch size for ADAN, we decided instead to use a conservative value further from its region of decreased performance at the expense of slower training. For reference, we also computed the execution time of PAF, which was much faster than both ADAN and Cycle-GAN (Fig. 4B, dashed blue line) as it has a closed form solution (23).

We next examined the effect of learning rates for each aligner. We first tested different values for the LR_G_, while fixing the ratio between LR_D_ and LR_G_ (for ADAN, LR_D_/LR_G_ = 0.5; for Cycle-GAN, LR_D_/LR_G_ = 10). As shown in Fig. 4C, ADAN achieved good performance when LR_G_ was set to 1e-5 and 1e-4 but did not work well if LR_G_ was set to 1e-3. Cycle-GAN maintained stable performance when LR_G_ was set to 1e-3 and 1e-4, and had a significant performance drop when LR_G_ was 1e-5 (1e-4: −0.064 ± 0.0062 (mean ± s.e.); 1e-5: −0.095 ± 0.0068; P ~ 0, Wilcoxon’s signed rank test), but still significantly better than ADAN with the same LR_G_ (Cycle-GAN: −0.095 ± 0.0068 (mean ± s.e.); ADAN: −0.15 ± 0.011; P ~ 0, Wilcoxon’s signed rank test). We then tested different ratios between LRD and LRG with LR_G_ fixed (LR_G_ = 1e-4 for both types of aligners). As Fig. 4D shows, ADAN could only be trained well when LRD was equal to or smaller than LRG. On the other hand, the performance of a Cycle-GAN based aligner remained stable for all tested LR_D_/LR_G_ values.

### GAN-based methods require very little training data for alignment

Aligners in practical iBCI applications must be fast to train and perhaps more importantly, require little training data. Here we investigated the aligner performance with limited training data. We trained ADAN, Cycle-GAN, and PAF to align the data on each day-k to four selected day-0s using randomly selected subsets of the full 120-trial training set from Monkey J. We then decoded EMGs from the aligned M1 signals on a fixed 40-trial held-out testing set using the day-0 decoder. As Fig. 5A shows, all three aligners improved the performance of day-0 decoders with 20 or fewer training trials. Performance increased as more training trials were included but started to plateau near 40 trials. When using only 10 trials, both ADAN and Cycle-GAN significantly outperformed PAF (Cycle-GAN: −0.19 ± 0.0076 (mean ± s.e.); ADAN: −0.21 ± 0.011; PAF: −0.26 ± 0.011; P ~ 0, Wilcoxon’s signed rank test), with Cycle-GAN significantly outperforming ADAN (P = 0.003, Wilcoxon’s signed rank test). It is also worth noting that ADAN and Cycle-GAN trained with only 20 trials significantly outperformed PAF trained with the full training set of 120 trials (Cycle-GAN trained with 20 trials: - 0.10 ± 0.0083 (mean ± s.e.); ADAN trained with 20 trials: −0.16 ± 0.0096; PAF trained with 120 trials: −0.20 ± 0.011; P ~ 0, Wilcoxon’s signed rank test) (Fig. 5B).

**Figure 5.**
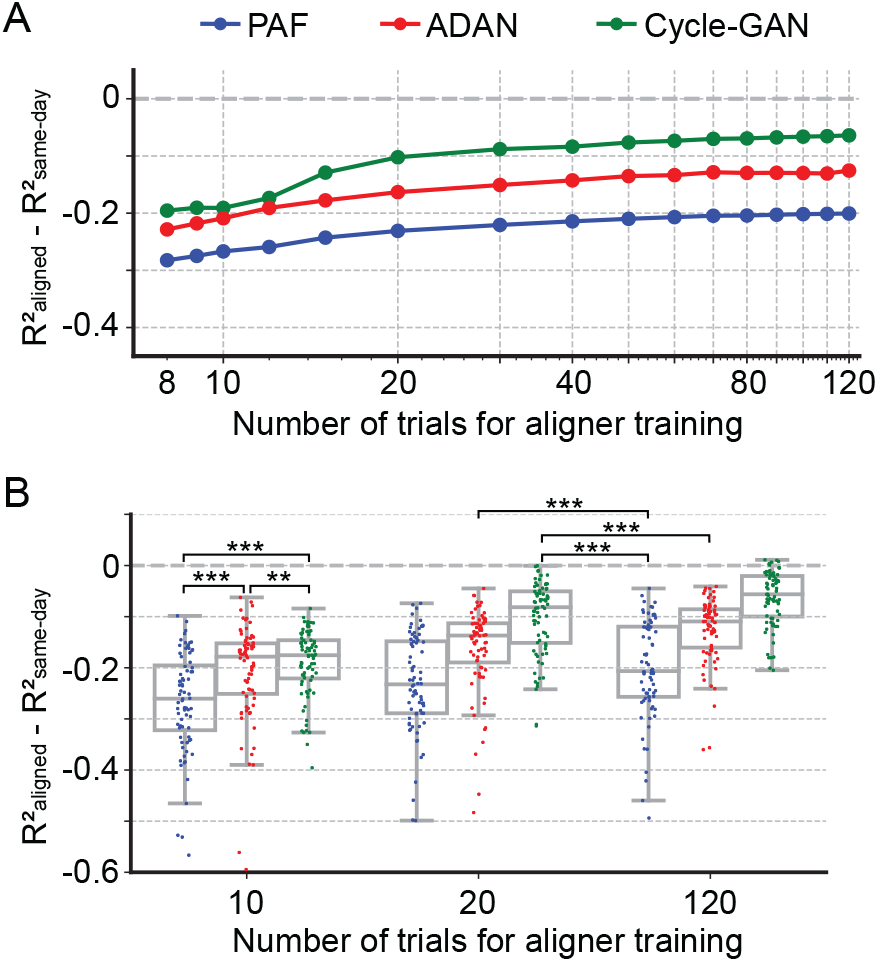
Cycle-GAN and ADAN need only a limited amount of data for training. (*A*) Effect of the number of trials used for training Cycle-GAN (green), ADAN (red) and PAF (blue) on the day-k decoding accuracy using 4 selected day-0 fixed decoders. All the aligners needed 20-40 trials to achieve a satisfactory performance, before reaching a plateau. The average prediction performance drop with respect to a daily-retrained decoder (R^2^_aligned_ - R^2^_same-day_) on all day-ks is shown for each tested value of training trials (x-axis is in log scale). When using 10 trials, both Cycle-GAN and ADAN significantly outperformed PAF (*B*, left boxplots). Moreover, both Cycle-GAN-based and ADAN aligners trained with 20 trials had significantly better performance than the PAF trained on all 120 trials (*B*, center and right boxplots). Single dots show the prediction performance drop on each day-k to the 4 selected day-0s with respect to a daily-retrained decoder. Boxplots show 25th, 50th and 75th percentiles of the R^2^ drop with the whiskers extending to the entire data spread, not including outliers. Asterisks indicate significance levels: *\*P* < 0.05, *\*\*P* < 0.01, \*\*\**P* < 0.001.

### Recovery of single-electrode activity patterns through alignment

Both ADAN and Cycle-GAN generate reconstructed versions of the aligned day-k single neuron signals, agnostic to downstream use. However, our objective of decoder stabilization does not require that the full distribution of day-0 responses be recovered: we need only recover signals that are relevant to the decoding dimension. Decoder performance alone therefore does not provide a complete picture of the quality of neural alignment. To more thoroughly investigate the extent to which distribution alignment introduces biases or artifacts in predicted neural responses, we first compared aligner predictions of single-neuron with those of their recorded day-0 analogs.

Because PAF operates directly on the low-dimensional neural manifold, it can only generate single-neuron responses in the aligned representation by projecting back out from the manifold. We found that a stabilized day-0 decoder that uses these reconstructed firing rates from the latent space of the PAF factors performs poorly (SI Appendix, Fig. S5C). In contrast to PAF, Cycle-GAN and ADAN each generate synthetic firing rates for the full neural population (although ADAN still relies on a low-dimensional manifold as an intermediate step). Therefore, we restricted our analysis of single-neuron properties on the outputs of ADAN and Cycle-GAN.

Specifically, we asked how response properties of the day-k “aligned neurons” differed from those of the neurons recorded on the same electrode on day-0. To do so, we examined the aligned neural representations generated by Cycle-GAN and ADAN, again using the 95-day isometric wrist task dataset of monkey J. We first compared the peri-event time histograms (PETHs) of firing rates before and after alignment, to determine how the aligners altered day-k neural activity at the level of single electrodes. The PETHs in Fig. 6A show three examples of the ways in which single-electrode signals may differ across days, and the change produced by alignment. Electrode E35 is an example of neuron drop-out, in which the activity captured on day-0 was not observed on day-95. The PETHs of aligned day-95 data matched those of day-0 for all force directions, demonstrating that on day 95 both ADAN and Cycle-GAN aligners synthesized appropriate neural activity (Fig. 6A). Second, E73 is an example of activity not present on day-0, but recorded on day-95. In this case, the day-95 activity was suppressed to match that on day-0. Finally, E60 is an example of consistent neural activity over the two days, which the aligners left unchanged.

**Figure 6.**
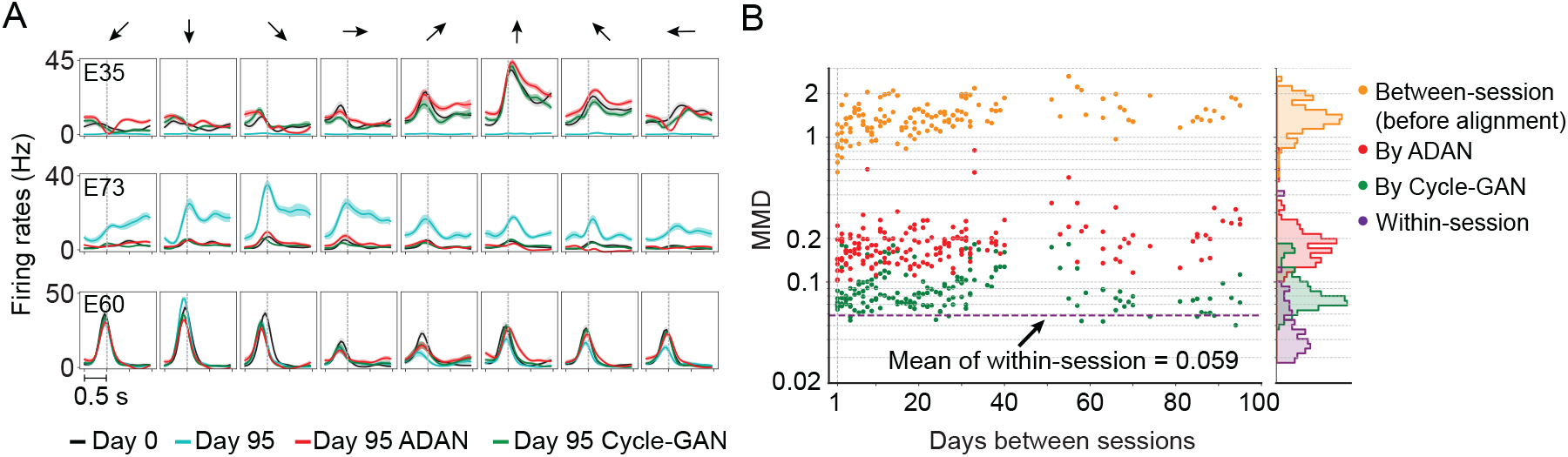
The changes of single-electrode and coordinated neural activity patterns after alignment. (*A*) The PETHs of the multiunit activity from three cortical electrodes (E35, E73, E60) before and after alignment. Each column corresponds to a target direction indicated by the arrows on the top. For each direction, mean (solid lines) and standard errors (shaded areas) are shown for 15 trials. The dashed vertical line in each subplot indicates the time of force onset. (*B*) Between-session MMDs for M1 signals before and after alignment, as well as the within-session MMDs. The main panel plots the between-session MMDs before (orange) and after alignment (red: by ADAN, green: by Cycle-GAN) for all pairs of sessions with different days apart, and the dashed purple line indicates the mean of the within-session MMD values. The side panel plots the histogram for each type of data. Note y-axis is in log scale.

We also examined the distributions of the recovered single-electrode activity by computing the Maximum Mean Discrepancy (MMD (24), see Methods) between all pairs of sessions (Fig. 6B). Before alignment, the between-day MMDs were significantly larger than the within-day MMDs (orange, between-day MMD: 1.42 ± 0.029 (mean ± s.e.); purple, within-day MMD: 0.059 ± 0.0054; P ~ 0, Wilcoxon’s rank sum test). After alignment, the between-day MMDs were substantially reduced by both Cycle-GAN and ADAN, becoming comparable to the within-day MMDs (ADAN: red, 0.19 ± 0.0065 (mean ± s.e.); Cycle-GAN: green, 0.091 ± 0.0024; within-day (purple, 0.059 ± 0.0054)). Cycle-GAN based aligners generally achieved a significantly lower between-day MMD than ADAN across the entire timespan (P ~ 0, Wilcoxon’s rank sum test).

### Recovery of neural manifolds from aligned representations

While Cycle-GAN works only with the full-dimensional neural recordings, ADAN, whose discriminator is essentially an autoencoder, computes a low-dimensional neural manifold from which it reconstructs the high-level signals it needs to align the high-level residuals. Consequently, we wanted to explore to what extent each method also altered the low-dimensional representations. We applied Principal Component Analysis (PCA) to the firing rates recorded for the 95-day isometric wrist task of monkey J on four selected day-0s and examined the trajectories of M1 neural activity within the neural subspaces defined by the principal components (PCs, see Methods). We then projected the firing rates of the remaining day-k’s onto the neural subspace defined by the corresponding day-0 PCs.

Generally, the day-k neural trajectories projected onto the top two day-0 PCs did not match those of day-0 (Fig. 7A). However, after alignment (3^rd^ and 4^th^ columns), the day-k trajectories closely resemble those of day-0.

**Figure 7.**
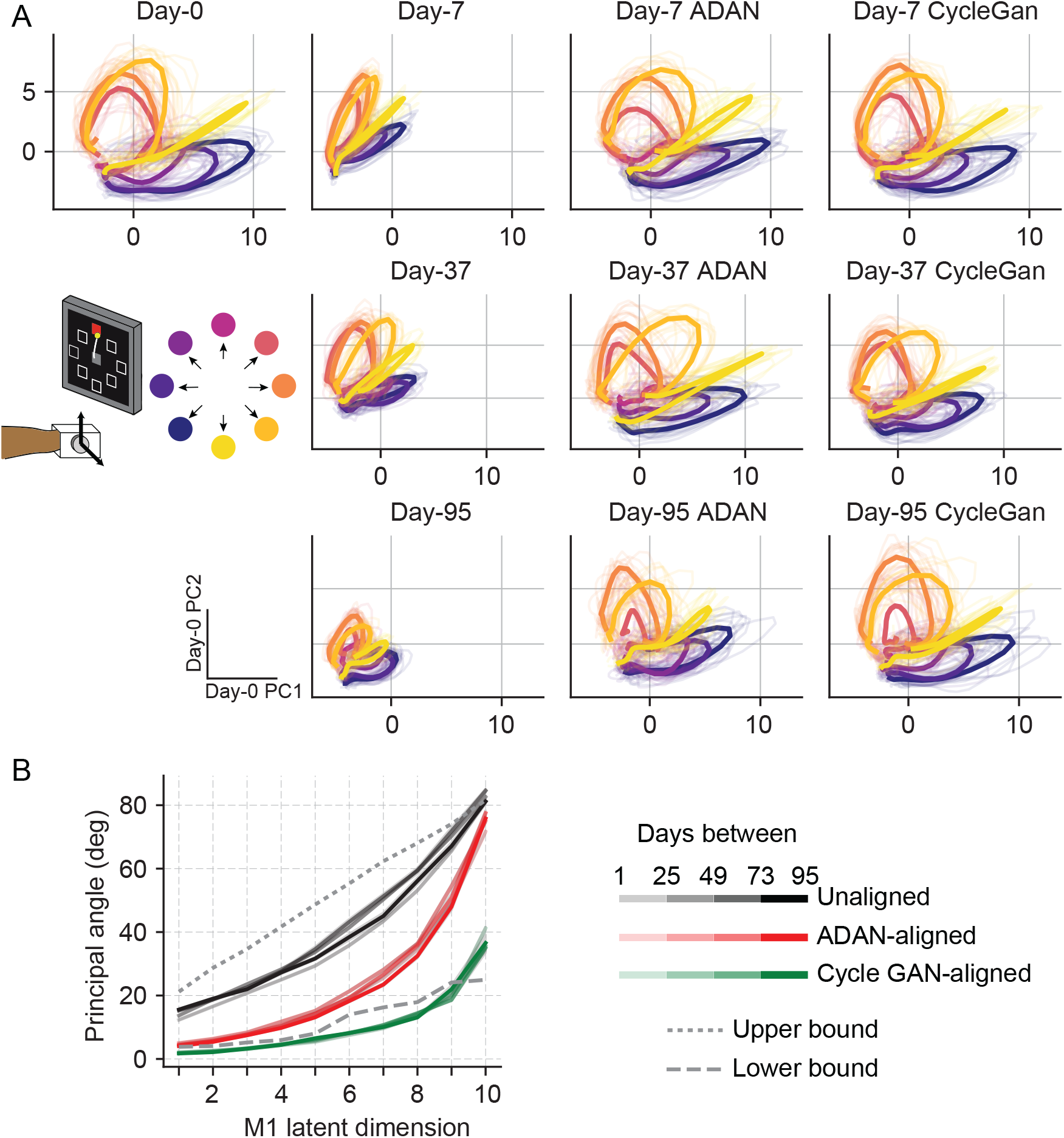
Neural manifold is stable over time after domain adaptation based neural alignment. (*A*) Representative latent trajectories when projecting unaligned / aligned neural activity onto the first two principal components (PCs) for the day-0 neural activity of monkey J during isometric wrist task. Top left corner: latent trajectories for day-0 firing rates, as the reference. 2^nd^ column: latent trajectories for unaligned firing rates on day-7 (top row), day-37 (center row) and day-95 (bottom row). 3^rd^ column and 4^th^ column: latent trajectories for firing rates aligned by ADAN (3^rd^ column) and Cycle-GAN (4^th^ column) on day-7, day-37, and day-95. Data were averaged over the first 16 trials for each target location and aligned to movement onset for visualization purposes. (*B*) First ten principal angles between the neural manifolds of day-0 and a given day-k for unaligned (black), aligned by ADAN (red) and aligned by Cycle-GAN (green). Upper bound was found by computing principal angles between surrogate subspaces with preserved statistics of day-0 and day-95 (0.01^st^ percentile is shown). Lower bound angles were found between subspaces relative to the first and second halves of day-0 neural recordings. Principal angle values were averaged across four different time intervals (relative to initial decoder training) indicated by the transparency of the line (lighter for days closer to day-0, darker for days further away from day-0).

Finally, to directly quantify the similarity between the neural manifolds of day-0 and an aligned day-k, we calculated the principal angles (25) between the neural subspaces for all sessions relative to the selected day-0 (see Methods). To interpret the magnitude of the overlap between a given pair of days, we compared the observed angle with an upper bound provided by the principal angles across random subspaces that preserved the covariance of the day-0 and day-95 neural data, using the method described in (26). We found a corresponding lower bound by splitting the neural recordings of day-0 in half and computing the principal angles between the subspaces of the two halves. We found that alignment with either Cycle-GAN or ADAN made the neural manifolds of any day-k substantially more similar to those of day-0, and in particular that after applying Cycle-GAN-based aligners, population subspaces highly overlapped (Fig. 7B).

## Discussion

We previously demonstrated the utility of a GAN-based method, ADAN, to “align” M1 data across time, thereby allowing a fixed iBCI decoder to be used for weeks without re-calibration, despite a gradual change in the neurons recorded over the same period (16). However, we had tested ADAN on a very limited dataset. Because GANs are notoriously sensitive to hyperparameter settings (18, 22, 27), it was unclear how robust ADAN would be in practice. Another promising method, PAF, had been tested primarily in terms of two monkeys’ online iBCI performance (9). We wished to compare both approaches directly, using a very diverse dataset including recordings from six monkeys and five tasks. We also compared a third approach based on a more recent GAN architecture, Cycle-GAN (19). Cycle-GAN has the potential advantage over ADAN that it reduces the search space of aligners by encouraging the learned transformation to be a bijection, which might help stabilize its performance. Moreover, unlike ADAN and PAF, the Cycle-GAN architecture does not require computation of a low-dimensional manifold underlying the neural population activity, allowing its straightforward use with spike-rate based decoders.

Both ADAN and Cycle-GAN achieved higher performance than PAF, but each method had tradeoffs. Although ADAN needed less time to train than Cycle-GAN, PAF was much faster to train than both GAN methods. But while slower, Cycle-GAN was easier to train than ADAN, in the sense that it was less sensitive to hyperparameter values. Overall, our work suggests that GAN-based alignment, and Cycle-GAN in particular, is a promising method for improving the stability of an iBCI over time.

### Comparison of GANs to other methods for iBCI stabilization

Other approaches to address iBCI decoder instability include supervised techniques that aim at stabilizing iBCI performance by recalibrating the decoder during ongoing iBCI control by relying on access to the task output variables (28–30), as well as unsupervised methods that do not require to re-estimate decoder parameters and only need neural data, with no provided task output variables or task labels (9, 16, 21, 31, 32). We restricted our comparison to GAN-based aligners and PAF for several reasons. First, both GANs and PAF are unsupervised methods. We argue that unsupervised methods are ideal for iBCI stabilization: because they do not require data labels, they should be simpler to implement in eventual clinical applications. Second, neither GANs nor PAF require time alignment of the data, which CCA does require. This flexibility allowed us to align the neural data for more complicated behaviors. For example, one task in this study was a random-target reaching task in which monkeys moved a cursor between targets as they appeared on screen; this task structure produces movements of random length and direction, with varied speed and duration. Despite this complexity, all three of the tested aligners could still achieve good performance. Importantly, though, we previously demonstrated that ADAN still achieves higher performance than both CCA (21) and KLDM (16) for the stereotyped isometric wrist task (16).

Although earlier attempts to achieve alignment via KLDM were only moderately successful, a recent approach using KLD to align neural latent dynamics identified using Latent Factor Analysis through Dynamical Systems (LFADS) (33, 34) was much more successful (32). In their evaluation, the resulting “Nonlinear Manifold Alignment with Dynamics” (NoMAD) also outperformed ADAN. However, there remain some differences between that study and ours. In part because of the considerable computational intensity of LFADS, Karpowicz et al. limited their analysis to a center-out reaching task with a kinematic decoder. Their train and test data were limited to a 750 ms window centered on movement onset, whereas we did not perform any temporal alignment (see Supplementary methods). Finally, they used a 20 ms bin size, which, if anything, would likely have decreased performance compared to our 50ms bins.

A very different approach to iBCI stabilization was proposed by Sussillo et al., who trained a decoder with a large dataset spanning many months, under the hypothesis that neural turnover allows neurons not only to disappear, but potentially also to reappear later (1). Although making the decoder robust to changes in the recorded neural populations, this approach has the inherent disadvantage of requiring the accumulation of a long stretch of historical data, which might be impractical for clinical use. For both Cycle-GAN and ADAN, there is no special requirement for the robustness of the day-0 decoder, and effective performance can be achieved with remarkably little data (Fig. 5).

### Sources of decoding error following cross-day alignment

In this study, we relied on offline estimates of decoder accuracy, as they allowed us to examine large amounts of previously collected data across many monkeys and tasks. Also, by literally taking the monkey out of the loop, we were able to examine the accuracy of the alignment and decoding processes without the added complication of the monkeys’ unknown and variable adaptation to the decoder. Although alignment by either ADAN or Cycle-GAN significantly improved the performance of a day-0 decoder on a given day-k, in most cases it did not attain the performance of a recalibrated decoder, especially at long time offsets between day-0 and day-k (Fig. 3B). One interesting potential cause of aligner performance drop is a change in the animal’s behavioral strategy across days. Because the limb is kinematically redundant, the same hand position can be achieved with different limb postures (e.g., wrist angle) and muscle activation patterns. Similarly, differing strategies might be adopted to grasp the power or pinch force transducers. Even within a single experimental session, an M1 decoder trained on one behavior often fails to perform well when tested on a different behavior. Similarly, unsupervised M1 alignment will not be able to compensate for changes in strategy if they shift EMG (or kinematic) signals outside the space of values observed during training of the original decoder. We find some evidence for such drift in some tasks (predominantly the key grasp, SI Appendix, Fig. S3C), as indicated by differences between within- and across-day MMD of the motor outputs. Such differences were small, but could not be neglected (SI Appendix, Fig. S2C, S3).

### Network training challenges

Training GANs is a challenging task, in part because the learning rates of generator and discriminator networks must be carefully balanced to allow the networks to be trained in tandem (18, 35). Many strategies have been proposed to improve the stability of learning and facilitate the convergence of GANs (18, 35–39). ADAN and Cycle-GAN incorporate several of those strategies. First, both networks include an L1 loss term in their objective function, a modification that has been found in practice to improve the stabilization of model training by encouraging sparseness of model weights (37). The networks also use a two-timescale update rule for generator and discriminator learning rates, which facilitates convergence of generator and discriminator to a balanced solution (40).

Correct optimization of GANs is also directly linked to proper tuning of the dynamics of learning during training (27, 41), which we investigated here in depth. Given the many GAN variants, there are still no comprehensive guidelines for a particular architecture (22). Consistent with this, we found that ADAN and Cycle-GAN differ substantially in their sensitivity to learning rate and batch size hyperparameters. Notably, ADAN exhibited poor generalization with larger batch sizes (like (42)), while Cycle-GAN worked well across all tested values (Fig. 4A). The ability to work with larger batch sizes gave Cycle-GAN several advantages over ADAN: its training was faster than ADAN (Fig. 4B) and it also enabled Cycle-GAN to maintain stable performance with higher learning rates (Fig. 4C and 4D, similar to the observations of (43)).

### Conclusions

In summary, we demonstrated the successful use of GANs for the stabilization of an iBCI, thereby overcoming the need for daily supervised re-calibration. Both approaches we tested (ADAN and Cycle-GAN) require remarkably little training data, making them practical for long-term iBCI clinical applications. Between the two approaches, Cycle-GAN achieved better performance which was less affected by inaccurate hyperparameter tuning; it is therefore our recommended method for future use. Notably, Cycle-GAN works directly with the unstable full-dimensional neural recordings, which further increases its performance and simplifies its implementation.

## Materials and Methods

### Subjects and behavior tasks

Six 9-10 kg adult male rhesus monkeys (Macaca mulatta) were used in this study. They were trained to sit in a primate chair and control a cursor on a screen in front of them using different behavioral apparatuses (SI Appendix, Fig. S1). Monkeys J and S were trained to perform an isometric wrist task, which required them to place their left hand inside a small box that was instrumented with a 6DOF load cell. They controlled the cursor on the screen by exerting forces within the box to track eight center-out targets. Monkeys P and G were trained to perform a grasping task, which required them to reach and grasp one of two force transducers located 30cm in front of their left shoulder. The transducers required either a cylindrical power grasp (monkey P) or a lateral key grasp (monkey G). Monkeys C and M were trained to perform a center-out (CO) reaching task while grasping the upright handle of a planar manipulandum, operated with the upper arm in a parasagittal plane. Monkey C performed the task with the right hand, monkey M with the left. Monkey M was also trained to perform a random-target (RT) task, reaching to a sequence of three targets presented in random locations on the screen to complete a single trial. More details about each behavior task can be found in Supplemental methods. All surgical and experimental procedures were approved by the Institutional Animal Care and Use Committee (IACUC) of Northwestern University, and are consistent with the Guide for the Care and Use of Laboratory Animals.

### Implants and data recordings

Depending on the task, we implanted a 96-channel Utah electrode array (Blackrock Neurotech, Inc.) in either the hand or arm representation area of the primary motor cortex (M1), contralateral to the arm being used for the task (see SI Appendix, Table S1). The implant site was pre-planned and finally determined during the surgery with reference to the sulcal patterns and the muscle contractions evoked by intraoperative surface cortical stimulation. For each of monkeys J, S, G, and P, we also implanted intramuscular leads in forearm and hand muscles of the arm used for the task in a separate procedure (see SI Appendix, Table S1). Electrode locations were verified during surgery by stimulating each lead.

M1 activity was recorded during task performance using a Cerebus system (Blackrock Neurotech, Inc.). The signals on each channel were digitalized, bandpass filtered (250 ~ 5000 Hz) and converted to spike times based on threshold crossings. The threshold was set with respect to the root-mean square (RMS) activity on each channel and kept consistent across different recording sessions (monkeys J, C and M: −5.5 x RMS; monkey S: −6.25 x RMS; monkey P: −4.75 x RMS; monkey G: −5.25 x RMS). The time stamp and a 1.6 ms snippet of each spike surrounding the time of threshold crossing were recorded. For all analyses in this study, we used multiunit threshold crossings on each channel instead of discriminating well isolated single units. We applied a Gaussian kernel (S.D. = 100 ms) to the spike counts in 50 ms, non-overlapping bins to obtain a smoothed estimate of firing rate as function of time for each channel.

The EMG signals were differentially amplified, band-pass filtered (4-pole, 50 ~ 500 Hz) and sampled at 2000 Hz. The EMGs were subsequently digitally rectified and low-pass filtered (4-pole, 10 Hz, Butterworth) and subsampled to 20 Hz. EMG channels with substantial noise were not included in the analyses, and data points of each channel were clipped to be no larger than the mean plus 6 times the S.D. of that channel. Within each recording session, we removed the baseline of each EMG channel by subtracting the 2nd percentile of the amplitudes and normalized each channel to the 90th percentile. For monkeys C and M, we recorded the positions of the endpoint of the reach manipulandum at a sampling frequency of 1000 Hz using encoders in the two joints of the manipulandum.

### iBCI day-0 decoder

The day-0 decoder was a Wiener filter of the type that we have used in several previous studies (6, 44). The filter was fit using linear regression to predict the motor outputs (either EMG or hand velocity) at time *t* given neural responses from time *t* to time *t -* T, where we set T = 4 (200 ms) for all decoders used in this study. As the aligners being tested worked with either low-dimensional manifolds or the full neural population, and required the associated day-0 decoders to be compatible, we implemented different day-0 decoders to match the outputs of the aligners. For Cycle-GAN, we trained a Wiener filter using the full-dimensional neural firing rates recorded on day-0. For ADAN and PAF, we performed dimensionality reduction (ADAN: autoencoder, PAF: Factor Analysis; dimensionality = 10 for both) to find a low-dimensional latent space, and trained the decoder using the projections of the neural signals into this latent space. The Wiener filters were trained using the day-0 data with 4-fold cross validation, and the filter corresponding to the fold with the best R^2^ was selected as the fixed day-0 decoder. The parameters for the dimensionality reduction procedures and the Wiener filter from the day-0 data were kept fixed for decoding on subsequent days.

### iBCI aligners

#### Adversarial domain adaptation network (ADAN)

We adhered to the main architecture and the training procedures of the ADAN as described in (16). Briefly, we first find a nonlinear latent space by jointly training an autoencoder and a long shortterm memory (LSTM) neural network-based iBCI decoder using day-0 data. (Note that this LSTM based decoder is only used for latent space discovery, not the later decoding stage that is used for performance evaluation (see “ADAN day-0 training” in supplementary methods for full details)). We then construct an adversarial aligner comprised of a distribution alignment module (generator network G) and a discriminator network D (SI Appendix, Fig. S7A), where G is a shallow feedforward neural network, and D is an autoencoder with the same architecture as that used for the day-0 latent space discovery. During training of the aligner, G is fed with day-k neural firing rates and applies a nonlinear transform over these data to match them to the day-0 neuron response distributions. The output of G, and the true day-0 neural firing rates are then passed to D, which passes both inputs through the autoencoder: namely, it projects each signal into the latent space and then reconstructs it. The distributions of the residuals between the autoencoder inputs and the reconstructions are computed for both the generator output and the true day-0 data, and a lower bound to the Wasserstein distance is used to measure the dissimilarity between the two distributions. The goal of adversarial learning is to find a discriminator D that maximizes the dissimilarity between responses of D to true day-0 firing rates and to outputs of G, while also finding a generator G that minimizes the dissimilarity between true day-0 firing rates and the outputs of G; this objective is called the adversarial loss. When the training is completed, G will have been trained to “align” the neural firing rates on day-k with those on day-0. For a full description of the ADAN architecture and its training strategy, please refer to the supplementary methods and (16).

#### Cycle-GAN

The Cycle-GAN aligner is based on the structure proposed in (19). Unlike ADAN, it converts the full-dimensional neural firing rates collected on day-k into a form resembling those collected on day-0, with no dimensionality reduction. Cycle-GAN consists of two feedforward generator neural networks (G1 and G2) and two discriminator networks (D1 and D2, see SI Appendix, Fig. S7B). These form two pairs of adversarial networks: G1 maps data from the day-k domain to the day-0 domain, while D1 aims to distinguish between the day-0 samples and the output of G1. And in parallel, G2 maps data in the day-0 domain to the day-k domain, while D2 distinguishes day-k data from output of G2. In contrast to ADAN, the cycle-GAN discriminator networks operate directly on neural responses, rather than the residuals between low-dimensional and full-dimensional responses.

The objective function for network training has two major terms. The first is an adversarial loss, defined for both generator-discriminator pairs (G1 + D1 and G2 + D2) as in ADAN. The second term is known as the cycle-consistency loss, which pushes the mappings G1 and G2 to become inverses of each other: that is, a sample from one specific domain should be recovered to its original form after going through the cycle composed of the two mappings. As argued by Zhu et al, the introduction of the cycle-consistency loss regularizes the learning of the mapping functions, thereby reducing the search space. In (SI Appendix, Fig. S7B) the purple arrows through G1 and G2 reflect the transformation of each sample from the day-k domain into the day-0 domain by G1, followed by the recovery from the day-0 domain into the day-k domain by G2. Likewise, the orange arrows through G2 and G1 reflect a transformation from the day-0 domain to the day-k domain and back to the day-0 domain. Further details about the Cycle-GAN based aligner are provided in the supplementary methods.

#### GAN training and architecture

Both ADAN and Cycle-GAN were trained using the ADAM optimizer (45) with a 4-fold cross validation. We used 400 training epochs and reported the alignment result that produced the best decoder performance on a held-out validation set of trials. In addition to the learning hyperparameters explored in the Results section, we examined several different architectures for the aligner neural network of both ADAN and Cycle-GAN (varying the number of layers and neurons per layer), and replaced the least absolute deviations (L1) for both the adversarial and cycle-consistency loss with the least square error (L2) (46). None of the manipulations substantially improved performance.

#### Procrustes alignment of factors (PAF)

We compared ADAN and Cycle-GAN aligners with a manifold-based stabilization method proposed by (9), the Procrustes Alignment of Factors (PAF, our term). PAF finds a low-dimensional manifold using Factor Analysis, then applies a Procrustes transformation to the neural manifold of day-0 to align it to that of day-k. The original application of PAF additionally removes electrodes identified as “unstable” and unlikely to contribute to alignment; these are defined as electrodes on day-k that have changed the most with respect to the day-0 manifold, and are removed iteratively until a criterion is met. As shown in (SI Appendix, Fig. S8), we found that alignment performance did not degrade with the number of included electrodes, so we decided to omit this stability criterion and use all recorded electrodes for all the datasets. As for the GAN aligners, we trained and tested PAF using a Wiener filter and 4-fold cross validation.

### Performance measures

#### Decoder accuracy

To evaluate the performance of decoders mapping M1 neural recordings to motor outputs (either EMG or hand velocity), we used the coefficient of determination (R^2^). The R^2^ indicates the proportion of variation of the actual motor output that was predicted by the iBCI decoder; this approach is common in evaluation of iBCI systems (47). As the motor outputs being decoded are multi-dimensional (7 dimensions for EMG, 2 dimensions for hand velocity), we computed a multivariate R^2^ in which, after computing the R^2^ for all the single dimensions, we take a weighted average across dimensions, with weights determined by the variance of each dimension. This was implemented using the “r2_score” function of the scikit-learn python package with “variance weighted” for the “multioutput” parameter (48).

#### Maximum mean discrepancy (MMD)

We used maximum mean discrepancy (MMD) in two contexts. First, we used MMD to evaluate the similarity between the distribution of the aligned day-k neural activity and the day-0 neural activity, as a way to examine the alignment performance (Fig. 6). MMD provides a measure of distance between two multivariate distributions, based on the distances between the mean embeddings of samples drawn from each distribution in a reproducing kernel Hilbert space (24). MMD is symmetric in the two distributions and equals zero if and only if the two distributions are the same. To select our kernel, we followed a technique that has been proved feasible for optimizing kernel choice (49): specifically we employed a family of four Gaussian kernels with width between 5 Hz and 50 Hz. To define a “smallest possible” MMD between aligned day-k and day-0 distributions, we divided neural signals recorded on the same day into non-overlapping folds, and computed MMD between them; we call this the “within-session MMD” in Fig. 6.

We also use the MMD to quantify the similarity of the distributions of neural activity or motor outputs between pairs of separate recording sessions for each dataset, as a way to quantify the recordings instabilities (SI Appendix, Fig. S2C, Fig. S3). For a pair of sessions, we divided each of them into 4 non-overlapping folds, and computed the MMD between each fold and its counterpart in the other session, then reported the mean value across folds. We also computed the “within-session MMD” for neural activity / motor outputs for each session, using the same way described above.

#### Principal angles

To evaluate the similarity between neural manifolds of day-0 and day-k before and after alignment, we used principal angles (25). Principal angles provide a metric to quantify the alignment of two subspaces embedded in a higher-dimensional space: principal angles approach zero when two manifolds have similar orientation, and approach 90 degrees when manifolds are nearly orthogonal. By construction, principal angles are ordered from smallest to largest. To compute the principal angle between datasets, we first computed the low-dimensional neural manifolds by applying PCA to the *C*-dimensional day-0 neural firing rates. Then, we projected the day-0 and day-k firing rates (both before and after alignment) of all *C* units into the embedding space described by the *C* day-0 PCs, to obtain the representative latent trajectories shown in Fig. 7A. We finally computed the principal angles between the day-0 and day-k subspaces for the first ten latent dimensions using the “subspace_angles” function of the SciPy python package (50).

To assess whether the angles after neural alignment were significantly small, we compared them to an upper bound provided by the angle between two surrogate subspaces, using the strategy described in (26). Briefly, we generated 10,000 random pairs of day-0-like and day-95-like subspaces in which we shuffled the timing of spikes within each neuron, destroying correlation structure while preserving the statistics of neural firing rates within each day. We then computed the principal angles between each pair, and used the 0.1^th^ percentile of the principal angle distribution as the threshold below which angles could be considered smaller than expected by chance given firing rate statistics alone. We also defined a lower bound by computing the principal angle between two day-0 subspaces derived from splitting the day-0 neural recordings in half and applying a separate PCA model on each. If the alignment process is successful, we expect the neural manifolds of day-0 and day-k to have similar principal angles to those indicated by the lower bound.

## Acknowledgments

We thank Ali Farshchian, Sara Solla and Ege Altan for valuable discussions. We thank current and former members of the Miller Limb Lab, including Kevin Bodkin, Stephanie Naufel, Matthew Perich, and Christian Ethier, for their contributions to data collection. The work was supported in part by grants to L.E.M. (R01 NS053603, R01 NS074044).

## Author Contributions

X.M., F.R. and L.E.M. designed research; X.M. and F.R. performed research; X.M. and F.R. analyzed data; X.M., F.R., A.K. and L.E.M. wrote the paper; and E.J.P., A.K., and L.E.M. provided supervision.

## Competing Interest Statement

The authors declare no competing interest.

## Data Availability

Data files from the 95-day isometric wrist task dataset of monkey J and all code used for analysis will be made publicly available on Dryad and GitHub ahead of manuscript publication.

## Notes

### Competing Interest Statement

The authors have declared no competing interest.

