## Supplementary Information for "Using adversarial networks to extend brain computer interface decoding accuracy over time"

**This PDF file includes:**

Supplementary methods
Supplementary results
Tables S1
SI References
Figures S1 to S8

### Supplementary methods

**Behavior tasks**. The isometric wrist torque task requires the monkey to control the cursor on the screen by exerting forces on a small box placed around one of the hands. The box was padded to comfortably constrain the monkey's hand and minimize its movement within the box, and the forces were measured by a 6 DOF load cell (JR3 Inc., CA) aligned to the wrist joint. During the task, flexion/extension force moved the cursor right and left respectively, while force along the radial/ulnar deviation axis moved the cursor up and down. Each trial started with the appearance of a center target requiring the monkeys to hold for a random time (0.2 – 1.0 s), after which one of eight possible outer targets selected in a block-randomized fashion appeared, accompanied with an auditory go cue. The monkey was allowed to move the cursor to the target within 2.0 s and hold for 0.8 s to receive a liquid reward (Fig. S1A). For both decoding and alignment analyses, we only used the data within each single trial (from 'trial start' to 'trial end', Fig. S1A). We did not do any temporal alignment with the trials, so the lengths of the trials were different from each other.

In the grasping tasks, monkeys were required to reach and grasp a gadget placed under the screen with one hand. The gadget was a cylinder for monkey P facilitating a power grasp with the palm and the fingers, while a small rectangular cuboid for monkey G facilitating a key grasp with the thumb and the index finger. A pair of force sensitive resistors (FSRs) were attached on the sides of the gadgets to measure the grasping forces the monkeys applied. The sum and the difference of the FSR outputs were used to determine the position of the cursor on the vertical axis and the horizontal axis respectively. At the beginning of each trial the monkey was required to keep the hand resting on a touch pad for a random time (0.5 – 1.0 s). A successful holding triggered the onset of one of three possible rectangular targets on the screen and an auditory go cue. The monkey was required to place the cursor into the target and hold for 0.6 s by increasing and maintaining the grasping force applied on the gadget (Fig. S1B). For this task we extracted trials from 'gocue time' to 'trial end', as the monkeys' movements were quite random before the gocue.

The center-out (CO) reach task required the monkey to manipulate a customized planar manipulandum. At the beginning of each trial the monkey needed to move the hand to the center of the workspace. One of eight possible outer targets equally spaced in a circle was presented to the monkey after a random waiting period. The monkey needed to keep holding for a variable delay period until receiving an auditory go cue. To receive a liquid reward, the monkey was required to reach the outer target within 1.0 s and hold within the target for 0.5 s (Fig. S1C). For this task we extracted trials from 'gocue time' to 'trial end', since the monkeys kept static before the gocue.

Monkey M was trained to perform the random-target (RT) task using the same apparatus for the CO reach task. At the beginning of each trial the monkey also needed to move the hand to the center of the workspace. Three targets were then presented to the monkey sequentially, and the monkey was required to move the cursor into each of them within 2.0 s after viewing each target. The positions of these targets were randomly selected, thus the cursor trajectory for each trial presented a 'random-target' manner (Fig. S1D). For this task we extracted trials from 'trial start' to 'trial end'.

**iBCI day-0 decoders.** We used a Wiener filter (5) as the day-0 iBCI decoder:

$$\boldsymbol{y}(t) = \sum_{\tau=0}^{\mathrm{T}-1} \boldsymbol{\beta}(\tau)\boldsymbol{x}(t-\tau) \qquad (1)$$

where $\boldsymbol{y}(t)$ is a q-dimensional vector ($\mathrm{q}$ is 2 for hand velocity prediction and varied with the number of recorded EMGs for EMG prediction, see Table S1) representing the motor outputs to be predicted at time $t$, while $\boldsymbol{x}(t)$ is a p-dimensional vector for the inputs to the Wiener filter at time $t$, and $\boldsymbol{\beta}(\tau)$ is a $\mathrm{q} \times \mathrm{p}$ matrix corresponding to the filter parameters for time step $\tau$. For Cycle-GAN, $\boldsymbol{x}(t)$ is the full-dimensional neural firing rates, thus p equals to the number of the electrodes in the cortical array (denoted as C). For ADAN, $\boldsymbol{x}(t)$ is the projection of the neural firing rates in a nonlinear latent space found by an autoencoder (see next section for details). For PAF, $\boldsymbol{x}(t)$ is the projection of the neural firing rates in a linear latent space found by factor analysis. We set $\mathrm{p} = 10$ for both ADAN and PAF. We can also write Eq. (1) in matrix form:

$$\boldsymbol{Y} = \boldsymbol{X}\boldsymbol{B} \tag{2}$$

where $\boldsymbol{Y}$ is a $\mathrm{M} \times \mathrm{q}$ matrix for the motor outputs to be predicted with M being the number of samples, $\boldsymbol{X}$ is a $\mathrm{M} \times (\mathrm{T} \times \mathrm{p})$ matrix, and $\boldsymbol{B}$ is a $(\mathrm{T} \times \mathrm{p}) \times \mathrm{q}$ matrix for the regression coefficients to be estimated. We also added an additional bias term for both $\boldsymbol{X}$ and $\boldsymbol{B}$. $\boldsymbol{B}$ was determined by a ridge regression estimator:

$$\hat{\boldsymbol{B}} = (\boldsymbol{X}^{\mathrm{T}}\boldsymbol{X} + \lambda\boldsymbol{I})^{-1}\boldsymbol{X}^{\mathrm{T}}\boldsymbol{Y} \tag{3}$$

We chose a ridge regression to limit the risk of decoder overfitting by penalizing solutions with large regression coefficients with the regularization term $\lambda$. The value of $\lambda$ was chosen by sweeping a range of 20 values between 10 and $10^5$ on a logarithmic scale. We used a 4-fold cross validation to train the decoder for each aligner type and ultimately selected the model with the highest $R^2$ on the test set as the fixed day-0 decoder.

**ADAN day-0 training.** The day-0 wiener filter for ADAN was built from a nonlinear latent space estimated from day-0 neural firing rates using an autoencoder (AE) originally described in (1). The AE consists of an input layer, five hidden layers and an output layer. The input and the output layers have $\mathrm{C}$ units, while the hidden layers (from input to output) have 64, 32, 10, 32 and 64 units, respectively. Hence, the AE compresses the $\mathrm{C}$-dimensional neural firing rates into a 10-dimensional latent representation. The units in the layer and the output layers as well as those in the latent layer have linear activation functions, while units in the remaining hidden layers have a nonlinear one (ELU). The AE is trained to minimize the reconstruction error defined as the mean square error (MSE) between the input and the output data. When day-0 neural firing rates $\{\boldsymbol{x}\}$ are fed through the AE, the latent layer activity $\{\boldsymbol{l}\}$ and the corresponding reconstructions $\{\hat{\boldsymbol{x}}\}$ are obtained. The 10-dimensional latent activity $\{\boldsymbol{l}\}$ is then mapped onto the q-dimensional motor output vector through a long-short-term memory (LSTM, (6)):

$$\hat{\boldsymbol{y}} = \mathrm{LSTM}(\boldsymbol{l}) \tag{4}$$

where $\boldsymbol{y}$ is the actual motor output (either EMG or hand velocity) recorded at day-0 and $\hat{\boldsymbol{y}}$ is its prediction with the LSTM. The LSTM is designed with one layer and a number of units that equals the number of recorded EMGs (if the motor output is EMG) or two (if the motor output is hand velocity). The AE and the LSTM are simultaneously trained by minimizing a loss function that accounts for both the MSE of the reconstruction of the firing rates ( $\mathcal{L}(\mathrm{AE})$ ) and the MSE of the motor output predictions ( $\mathcal{L}(\mathrm{LSTM})$ ):

$$\mathcal{L} = \lambda\mathcal{L}(\mathrm{AE}) + \mathcal{L}(\mathrm{LSTM}) = \frac{1}{\mathrm{M}}\sum_{i=1}^{\mathrm{M}}(\lambda \, || \, \hat{\boldsymbol{x}} - \boldsymbol{x} \, ||^2 + || \, \hat{\boldsymbol{y}} - \boldsymbol{y} \, ||^2) \tag{5}$$

where M is the total number of training samples. The weighting factor $\lambda$ equalizes the contribution of the two terms so that the learning algorithm does not prioritize one over the other. For each training epoch, $\lambda$ is updated as the ratio between the values of $\mathcal{L}(\mathrm{AE})$ and $\mathcal{L}(\mathrm{LSTM})$ at the end of the preceding epoch.

The simultaneous training of the AE and the LSTM allows extracting a low-dimensional space of neural activity constrained to include features related to movement intent. Such neural manifold is then used to train the Wiener filter used as the fixed day-0 decoder for this study. At each epoch of training, the current latent signal $\{\boldsymbol{l}\}$ was used as input for Eq. (3) to obtain a linear prediction of the actual motor output. We used 400 epochs of training and ultimately selected the parameters of the wiener filter at the epoch that had the best performance (in the $R^2$ sense) on the held-out test set.

**ADAN based aligner.** The discriminator $\mathrm{D}$ of ADAN is an autoencoder (Fig. S7A), and has the same architecture as that used to find the nonlinear latent space on day-0 (day-0 AE). The parameters of $\mathrm{D}$ ($\boldsymbol{\theta}_{\mathrm{D}}$) are initialized with the parameters of the day-0 AE. The generator G is a feedforward neural network with one hidden layer with $\mathrm{C}$ neurons (i.e., the number of the electrodes in the cortical array). The parameters of G ($\boldsymbol{\theta}_{\mathrm{G}}$) are initialized as identity matrices. We set a nonlinear activation function (exponential linear unit, ELU) for the hidden layer, and a linear one for the output layer.

Here we denote the day-0 neural firing rates as $\{\boldsymbol{x}_i\}_{i=1}^{\mathrm{M}}$ and the day-k neural firing rates as $\{\boldsymbol{z}_j\}_{j=1}^{\mathrm{N}}$, where both $\boldsymbol{x}_i$ and $\boldsymbol{z}_j$ are $\mathrm{C}$-dimensional vectors representing the neural firing rates from $\mathrm{C}$ electrodes at a given time bin, and $\mathrm{M}$ and $\mathrm{N}$ are the total number of samples for day-0 and day-k data respectively. Since at one time we fed the networks with S training samples as a batch, we can write a training batch from $\{\boldsymbol{x}\}$ or $\{\boldsymbol{z}\}$ in matrix form as $\boldsymbol{X}$ or $\boldsymbol{Z}$. During training, we fed $\boldsymbol{Z}$ to $\mathrm{G}$ and got $\mathrm{G}(\boldsymbol{Z})$ as the aligned day-k neural firing rates. At the same time, we fed $\mathrm{D}$ with both $\mathrm{G}(\boldsymbol{Z})$ and $\boldsymbol{X}$. As $\mathrm{D}$ is an autoencoder, it would produce the reconstructions of them from the latent space, which can be written as $\widehat{\mathrm{G}(\boldsymbol{Z})}$ and $\widehat{\boldsymbol{X}}$. Hence, we could get the residuals between the true data and these reconstructions by computing:

$$
\begin{aligned}
\boldsymbol{R}_{X} &= \boldsymbol{X} - \widehat{\boldsymbol{X}} \\
\boldsymbol{R}_{\mathrm{G}(\boldsymbol{Z})} &= \mathrm{G}(\boldsymbol{Z}) - \widehat{\mathrm{G}(\boldsymbol{Z})}
\end{aligned}
\tag{6}
$$

$\boldsymbol{R}_X$ and $\boldsymbol{R}_{\mathrm{G}(\boldsymbol{Z})}$ are both $\mathrm{S}\times\mathrm{C}$ matrices. We then computed the scalar reconstruction losses as the $L_1$ norm of each column of $\boldsymbol{R}_X$ and $\boldsymbol{R}_{\mathrm{G}(\boldsymbol{Z})}$. Let $\rho(\boldsymbol{R}_X)$ and $\rho(\boldsymbol{R}_{\mathrm{G}(\boldsymbol{Z})})$ represent the distributions of these scalar losses, and let $\mu(\boldsymbol{R}_X)$ and $\mu(\boldsymbol{R}_{\mathrm{G}(\boldsymbol{Z})})$ be the corresponding means of $\rho(\boldsymbol{R}_X)$ and $\rho(\boldsymbol{R}_{\mathrm{G}(\boldsymbol{Z})})$. We measured the dissimilarity between $\rho(\boldsymbol{R}_X)$ and $\rho(\boldsymbol{R}_{\mathrm{G}(\boldsymbol{Z})})$ by a lower bound to the Wasserstein distance (2), which is given by the absolute value of the difference between $\mu(\boldsymbol{R}_X)$ and $\mu(\boldsymbol{R}_{\mathrm{G}(\boldsymbol{Z})})$: $\mathrm{W}(\rho(\boldsymbol{R}_X),\rho(\boldsymbol{R}_{\mathrm{G}(\boldsymbol{Z})})) \geq |\mu(\boldsymbol{R}_X)-\mu(\boldsymbol{R}_{\mathrm{G}(\boldsymbol{Z})})|$. The parameters of the generator ($\boldsymbol{\theta}_{\mathrm{G}}$) and discriminator ($\boldsymbol{\theta}_{\mathrm{D}}$) are updated via batch gradient descent by minimizing their corresponding cost functions:

$$
\begin{aligned}
\mathcal{L}(\mathrm{D}) &= \mu(\boldsymbol{R}_X) - \mu(\boldsymbol{R}_{\mathrm{G}(\boldsymbol{Z})}) \\
\mathcal{L}(\mathrm{G}) &= \mu(\boldsymbol{R}_{\mathrm{G}(\boldsymbol{Z})})
\end{aligned}
\tag{7}
$$

For each epoch of training, $\mathcal{L}(\mathrm{G})$ is first minimized and followed by $\mathcal{L}(\mathrm{D})$. Minimizing $\mathcal{L}(\mathrm{G})$ implies bringing the output of the generator (*i.e.*, the aligned day-k neural data, $\mathrm{G}(\boldsymbol{Z})$) close to the day-0 data $\boldsymbol{X}$. When $\mathrm{G}(\boldsymbol{Z})$ is fed through $\mathrm{D}$, residuals with mean $\mu_Z$ are obtained. Since $\mathrm{D}$ is initialized with the day-0 AE weights, $\mu_Z$ can be reduced if $\theta_{\mathrm{G}}$ are updated to appropriately modify $\mathrm{G}(\boldsymbol{Z})$ and make it resemble $\boldsymbol{X}$. When $\mathcal{L}(\mathrm{G})$ is minimized, the gradients flow through both $\mathrm{D}$ and $\mathrm{G}$, but only the parameters $\boldsymbol{\theta}_{\mathrm{G}}$ are updated at this stage.

While G is trying to decrease $\mu_{\boldsymbol{Z}}$ , D is working as an adversary. Minimizing $\mathcal{L}(\mathrm{D})$ implies maximizing the difference between $\mu(\boldsymbol{R}_{\boldsymbol{X}})$ and $\mu(\boldsymbol{R}_{\mathrm{G}(\boldsymbol{Z})})$ (*i.e.*, their Wasserstein distance W). Again, since D is initialized with the day-0 AE weights (and the generator is an identity matrix when training begins), the residuals of the day-k data will be greater than those of the day-0 data, hence $(\mu_{\boldsymbol{Z}} > \mu_{\boldsymbol{X}})$. Thus, if $\boldsymbol{\theta}_{\mathrm{D}}$ are updated to maximize $(\mu_{\boldsymbol{Z}} - \mu_{\boldsymbol{X}})$, or equivalently minimize $(\mu_{\boldsymbol{X}} - \mu_{\boldsymbol{Z}})$, this relation is maintained during training. Since scalar residuals and their means are always nonnegative, maximization of W is achieved by decreasing $\mu_{\boldsymbol{X}}$ while increasing $\mu_{\boldsymbol{Z}}$ . The adversarial mechanism between G and D ensures that the neural alignment is achieved in an unsupervised manner.

**Cycle-GAN based aligner.** The Cycle-GAN generators, $\mathrm{G}_1$ and $\mathrm{G}_2$ are both shallow feedforward neural networks with one hidden layer with C neurons. We set a nonlinear activation function (ELU) for the hidden layer, and a linear one for the output layer. The discriminators, $\mathrm{D}_1$ and $\mathrm{D}_2$ are also shallow feedforward neural networks with one hidden layer. The input layer and the hidden layer both have C neurons, while the output layer has 1 neuron, as the output is a class label indicating which distribution the input sample belongs to. Same as $\mathrm{G}_1$ and $\mathrm{G}_2$, the hidden layer of $\mathrm{D}_1$ and $\mathrm{D}_2$ uses a nonlinear activation function (ELU), and the output layer uses a linear one. The layer weights of each network were initialized through Xavier initialization.

As shown in Fig. S7B, we fed the day-k neural firing rates $\boldsymbol{Z}$ to $\mathrm{G}_1$ to get the aligned day-k neural firing rates ($\mathrm{G}_1(\boldsymbol{Z})$), and the day-0 neural firing rates $\boldsymbol{X}$ to $\mathrm{G}_2$ to convert data in the day-0 domain back into the day-k domain ($\mathrm{G}_2(\boldsymbol{X})$). Meanwhile, the discriminator $\mathrm{D}_1$ was fed with $\boldsymbol{X}$ and ($\mathrm{G}_1(\boldsymbol{Z})$) to distinguish between the 'real and the 'fake' day-0 data, while $\mathrm{D}_2$ was fed with $\boldsymbol{Z}$ and ($\mathrm{G}_2(\boldsymbol{X})$) to distinguish between the 'real' and the 'fake' day-k data. Specifically, the discriminators would assign each sample a class label to tell if it belonged to the C-dimensional distribution of the real data ( $\rho(\boldsymbol{X})$ or $\rho(\boldsymbol{Z})$ ) or from the distribution of the fake data generated by $\mathrm{G}_1$ or $\mathrm{G}_2$.

For the network training, we expected $\mathrm{G}_1$ and $\mathrm{G}_2$ to generate more convincing samples, while $\mathrm{D}_1$ and $\mathrm{D}_2$ to be more perceptive to better discriminate between the true and the fake samples. The performances of the networks in such contest could be quantified by adversarial losses. As with ADAN, here we adopted the mean absolute error (MAE), or L1 loss, as the adversarial loss function. For $\mathrm{G}_1$ and $\mathrm{D}_1$, the adversarial loss can be expressed as follows:

$$\begin{aligned}\mathcal{L}_{adv}(\mathrm{D}_1) &= \mathrm{E}_{\boldsymbol{X}\sim \mathrm{p}_{\mathrm{data}}(\boldsymbol{X})}[\| \mathrm{D}_1(\boldsymbol{X}) - b \|_1] + \mathrm{E}_{\boldsymbol{Z}\sim \mathrm{p}_{\mathrm{data}}(\boldsymbol{Z})}[\| \mathrm{D}_1(\mathrm{G}_1(\boldsymbol{Z})) - a \|_1] \\ \mathcal{L}_{adv}(\mathrm{G}_1) &= \mathrm{E}_{\boldsymbol{Z}\sim \mathrm{p}_{\mathrm{data}}(\boldsymbol{Z})}[\| \mathrm{D}_1(\mathrm{G}_1(\boldsymbol{Z})) - c \|_1]\end{aligned} \tag{8}$$

where $a$ is the label for the fake neural firing rates, $b$ is the label for the real neural firing rates, and $c$ is the value that $\mathrm{G}_1$ wants $\mathrm{D}_1$ to believe for fake neural firing rates. Typically, we can set $a = 0$, and $b = c = 1$. For $\mathrm{D}_2$ and $\mathrm{G}_2$, the adversarial loss $\mathcal{L}_{adv}(\mathrm{D}_2)$ and $\mathcal{L}_{adv}(\mathrm{G}_2)$ have a similar form:

$$\begin{aligned}\mathcal{L}_{adv}(\mathrm{D}_2) &= \mathrm{E}_{\boldsymbol{Z}\sim \mathrm{p}_{\mathrm{data}}(\boldsymbol{Z})}[\| \mathrm{D}_2(\boldsymbol{Z}) - b \|_1] + \mathrm{E}_{\boldsymbol{X}\sim \mathrm{p}_{\mathrm{data}}(\boldsymbol{X})}[\| \mathrm{D}_2(\mathrm{G}_2(\boldsymbol{X})) - a \|_1] \\ \mathcal{L}_{adv}(\mathrm{G}_2) &= \mathrm{E}_{\boldsymbol{X}\sim \mathrm{p}_{\mathrm{data}}(\boldsymbol{X})}[\| \mathrm{D}_2(\mathrm{G}_2(\boldsymbol{X})) - c \|_1]\end{aligned} \tag{9}$$

The core idea of Cycle-GAN is to make the learned mapping functions cycle-consistent so as to reduce the space of possible mapping functions. As shown in Fig. S7B, the two highlighted cycles should be able to bring the corresponding data back to the original domain, for example, the distribution of the recovered day-k neural firing rates $\mathrm{G}_2(\mathrm{G}_1(\boldsymbol{Z}))$ should be similar to the distribution of the real day-k neural firing rates $\boldsymbol{Z}$. Therefore, we define the cycle consistency loss as follows:

$$\mathcal{L}_{cyc}(\mathrm{G}_1, \mathrm{G}_2) = \mathrm{E}_{\boldsymbol{X}\sim \mathrm{p}(\boldsymbol{X})}[\| \mathrm{G}_1(\mathrm{G}_2(\boldsymbol{X})) - \boldsymbol{X} \|_1] + \mathrm{E}_{\boldsymbol{Z}\sim \mathrm{p}(\boldsymbol{Z})}[\| \mathrm{G}_2(\mathrm{G}_1(\boldsymbol{Z})) - \boldsymbol{Z} \|_1] \tag{10}$$

Note here we also applied the L1 loss.

Taken together, the full loss function is written as:

$$\mathcal{L}(G_1, G_2, D_1, D_2) = \mathcal{L}_{adv}(D_1) + \mathcal{L}_{adv}(G_1) + \mathcal{L}_{adv}(D_2) + \mathcal{L}_{adv}(G_2) + \mathcal{L}_{cyc}(G_1, G_2) \quad (11)$$

and the training process is to solve this min-max optimization problem:

$$G_1^*, G_2^*, D_1^*, D_2^* = \arg\min_{G_1, G_2} \max_{D_1, D_2} \mathcal{L}(G_1, G_2, D_1, D_2) \quad (12)$$

**Supplementary results**

**Unstable neural recordings underlying stable motor outputs.** We evaluated the stability of the M1 neural activity as well as the motor outputs across time. The peri-event time histograms (PETHs) of M1 signals (Fig. S2A, top panel) from monkey J, who was trained to perform the isometric wrist task, show that the neural activity picked by the implanted electrodes may change dramatically (E35, E73) or remain largely consistent (E95) over days. In contrast, the EMG patterns (Fig. S2A, bottom panel) from two muscles which are critical to the task (FCU, ECRl) remained stable even over 95 days. These observations can be well characterized by the probability distribution functions (PDFs): the PDFs for different days deviated from each other if the patterns of the signals changed, as shown in Fig. S2B. We then utilized maximum mean discrepancy (MMD) to measure the difference of the PDFs over 96-dimensional M1 neural signals between pairs of recording sessions. We also obtained the within-session MMDs by dividing each recording session into four non-overlapping folds and computing the MMD between the folds. As shown in Fig. S2C, the between-session MMDs for M1 signals were at least ten times greater than the mean of the within-session MMDs (0.059) even there was only one day apart between two sessions, and tended to be larger when there were more days apart. These results suggest the prevalence of the instabilities featured by the aforementioned E35 and E73 in the neural recordings, which caused the shift of the overall distributions of M1 signals across time. In contrast, when using MMD to measure the difference of the PDFs over all EMG channels between sessions (Fig. S2C, bottom panel), we found that all between-session MMDs were very small (<0.04). Such results suggest the motor outputs during the behavioral task were generally stable over time. Since the monkey was already well trained and proficient with the tasks before the whole data collection process began, the motor outputs would not change greatly across time as the behavior during task performing had become stable. However, there are also some factors that may slightly alter the actual or measured motor outputs across time, such as monkey's daily condition, noise levels of recordings, and drifts of the sensors on the behavioral apparatus. They could altogether contribute to the gradual increase of the cross-session MMDs of EMGs over time shown in Fig. S2C. We performed the same analyses for all monkeys / tasks and observed similar results (Fig. S3).

**Table S1.** Information for each dataset.

| Task | Monkey | Cortical implant site | Cortical implant date | Number of recording sessions | Range of days | Range since array implantation | Motor outputs being recorded * |
|---|---|---|---|---|---|---|---|
| Isometric wrist | J | Right M1 (hand area) | 2013-09-10 | 20 | 95 | 688 - 783 | EMGs from 7 muscles in left arm (ECU, FCU, ECRl, EDC, FCR, FDP, ECRb) |
| | S | Right M1 (hand area) | 2012-05-07 | 18 | 83 | 106 - 189 | EMGs from 8 muscles in left arm and hand (FDP1, FDP2, FCR, 1DI, FPB, EDC, MD, ECRl) |
| Grasping ** | G | Right M1 (hand area) | 2019-07-23 | 8 | 53 | 23 - 76 | EMGs from 8 muscles in left arm and hand (FCR, FDP, PT, FPB, 1DI, SUP, ECU, EDC) |
| | P | Right M1 (hand area) | 2021-05-19 | 9 | 51 | 14 - 65 | EMGs from 11 muscles in left arm and hand (APB, FPB, Lum, PT, FDS, FDP, 1DI, 4DI, EPL, ECRl, EDC) |
| Center-out reach | C | Left M1 (arm area) | 2016-08-09 | 12 | 38 | 49 - 87 | Hand velocities ($v_x$, $v_y$) |
| | M | Right M1 (arm area) | 2013-06-06 | 11 | 32 | 242 - 274 | Hand velocities ($v_x$, $v_y$) |
| Random-target reach | M | Right M1 (arm area) | 2013-06-06 | 11 | 79 | 184 - 263 | Hand velocities ($v_x$, $v_y$) |

* Abbreviations for the muscles: ECU (extensor carpi ulnaris), FCU (flexor carpi ulnaris), ECRl (extensor carpi radialis longus), EDC (extensor digitorum communis), FCR (flexor carpi radialis), FDP (flexor digitorum profundus), ECRb (extensor carpi radialis brevis), FCR (flexor carpi radialis), 1DI (first dorsal interosseous), FPB (flexor pollicis brevis), MD (opponens digiti minimi), PT (pronator), SUP (supinator), APB (abductor pollicis brevis), Lum (lumbrical), FDS (flexor digitorum superficialis), 4DI (fourth dorsal interosseous), EPL (extensor pollicis longus)

** Monkey G was trained to do key grasping, and Monkey P do power grasping

*** For grasping, center-out, and random-target reach we tested all the available recording sessions as day-0, while for the isometric wrist task we tested 9 out of the available 20 sessions for monkey J and 8 out of the available 18 for monkey S. Each day in a dataset other than the designated day-0 was treated as a day-k, whether it occurred before or after day-0.

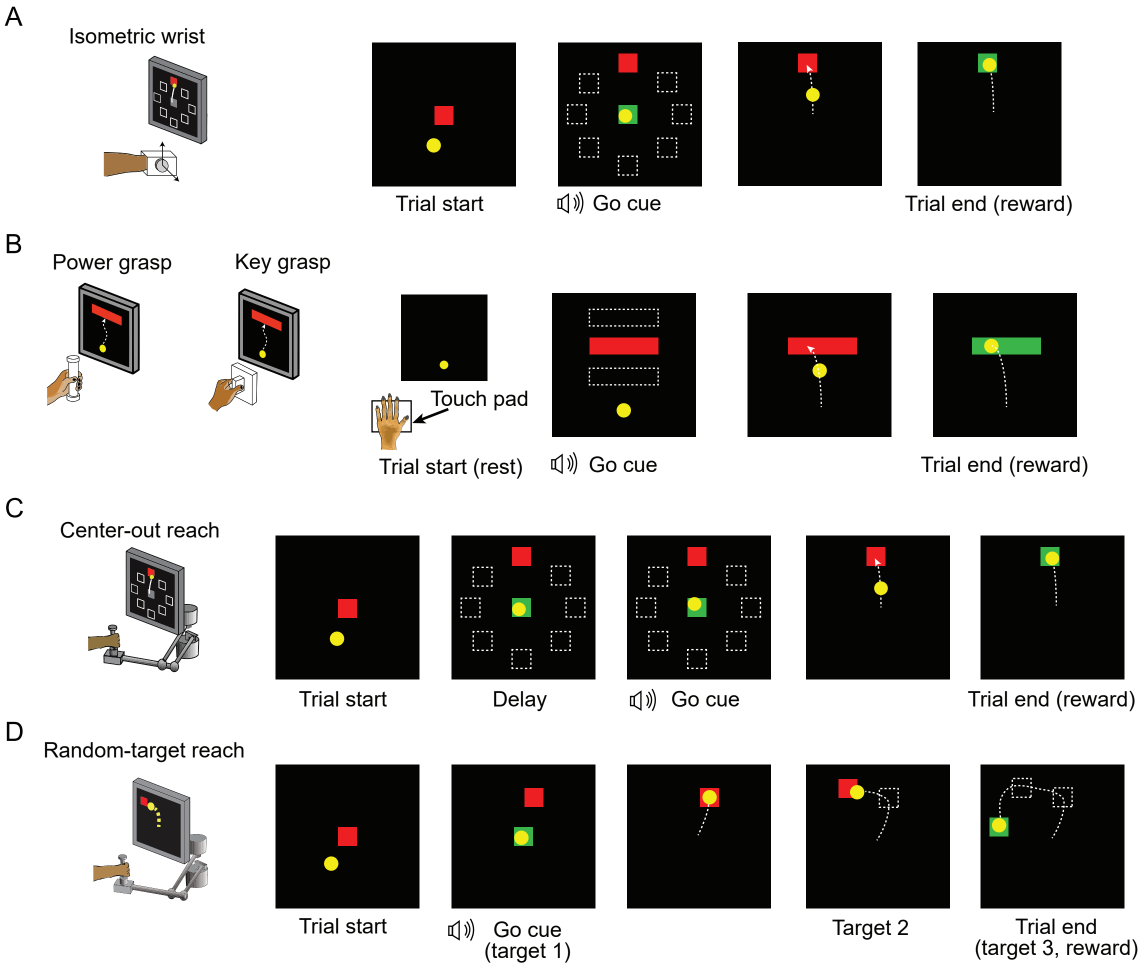


**Figure S1**. Behavior tasks. (*A*) The structure of the isometric wrist task. Each trial started with the appearance of a center target requiring the monkeys to hold for a random time (0.2 – 1.0 s), after which one of eight possible outer targets selected in a block-randomized fashion appeared, accompanied with an auditory go cue. The monkey was allowed to move the cursor to the target within 2.0 s and hold for 0.8 s to receive a liquid reward. (*B*) The structure of the grasping tasks. At the beginning of each trial the monkey was required to keep the hand resting on a touch pad for a random time (0.5 – 1.0 s). A successful holding triggered the onset of one of three possible rectangular targets on the screen and an auditory go cue. The monkey was required to place the cursor into the target and hold for 0.6 s by increasing and maintaining the grasping force applied on the gadget. (*C*) The structure of the center-out (CO) reach task. At the beginning of each trial the monkey needed to move the hand to the center of the workspace. One of eight possible outer targets equally spaced in a circle was presented to the monkey after a random waiting period. The monkey needed to keep holding for a variable delay period until receiving an auditory go cue. To receive a liquid reward, the monkey was required to reach the outer target within 1.0 s and hold within the target for 0.5 s. (*D*) The structure of the random-target (RT) reach task. At the beginning of each trial the monkey also needed to move the hand to the center of the workspace. Three targets were then presented to the monkey sequentially, and the monkey was required to move the cursor into each of them within 2.0 s after viewing each target. The positions of these targets were randomly selected, thus the cursor trajectory for each trial presented a 'random-target' manner.

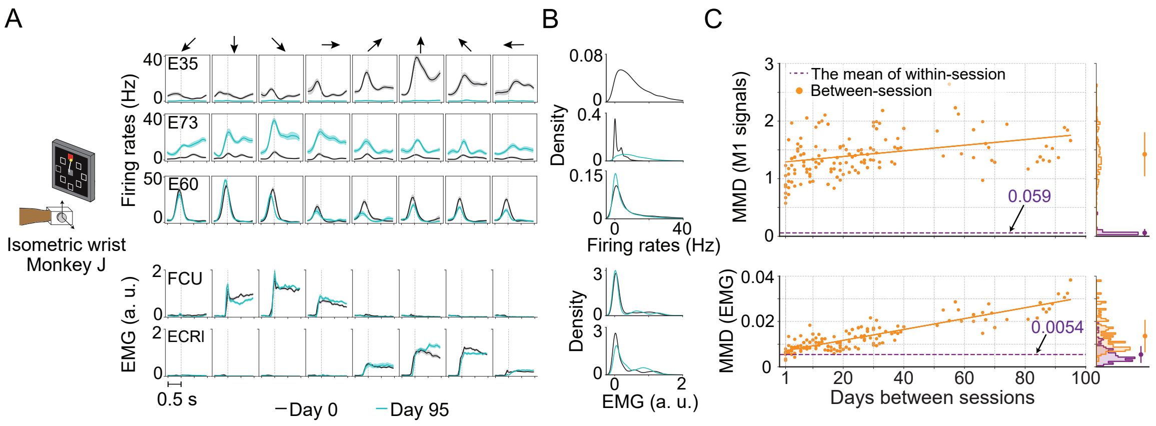


**Figure S2**. Unstable neural recordings underlying stable motor outputs. Data were from monkey J, who was trained to perform the isometric wrist task. (*A*) Peri-event time histograms (PETHs) for the multiunit activity from three cortical electrodes (E35, E73, E60) and the EMGs from two forearm muscles (flexor carpi ulnaris, FCU; extensor carpi radialis longus, ECRl) on day 0 and day 95. Each column corresponds to a target direction indicated by the arrows on the top. For each direction 15 trials were averaged to get the mean values (solid lines) and the standard errors (shaded area). The dashed vertical line in each subplot indicates the timing of force onset. (*B*) The distributions of the neural firing rates from E35, E73 and E60 and the EMGs from FCU and ECRl. The order of the subplots is consistent with (A). Note that for E35 the distribution for day-95 neural firing rates was omitted, since all values were close to 0. (*C*) The within-session and between-session MMDs for M1 signals (top panel) and EMGs (bottom panel). In each panel the solid orange line shows a linear fit for all between-session MMDs, the dashed purple line indicates the mean of all within-day MMD values. The histograms for within-session and between-session MMDs were plotted on the right side of each panel, and the mean (solid dots) and standard deviation (solid lines) were shown.

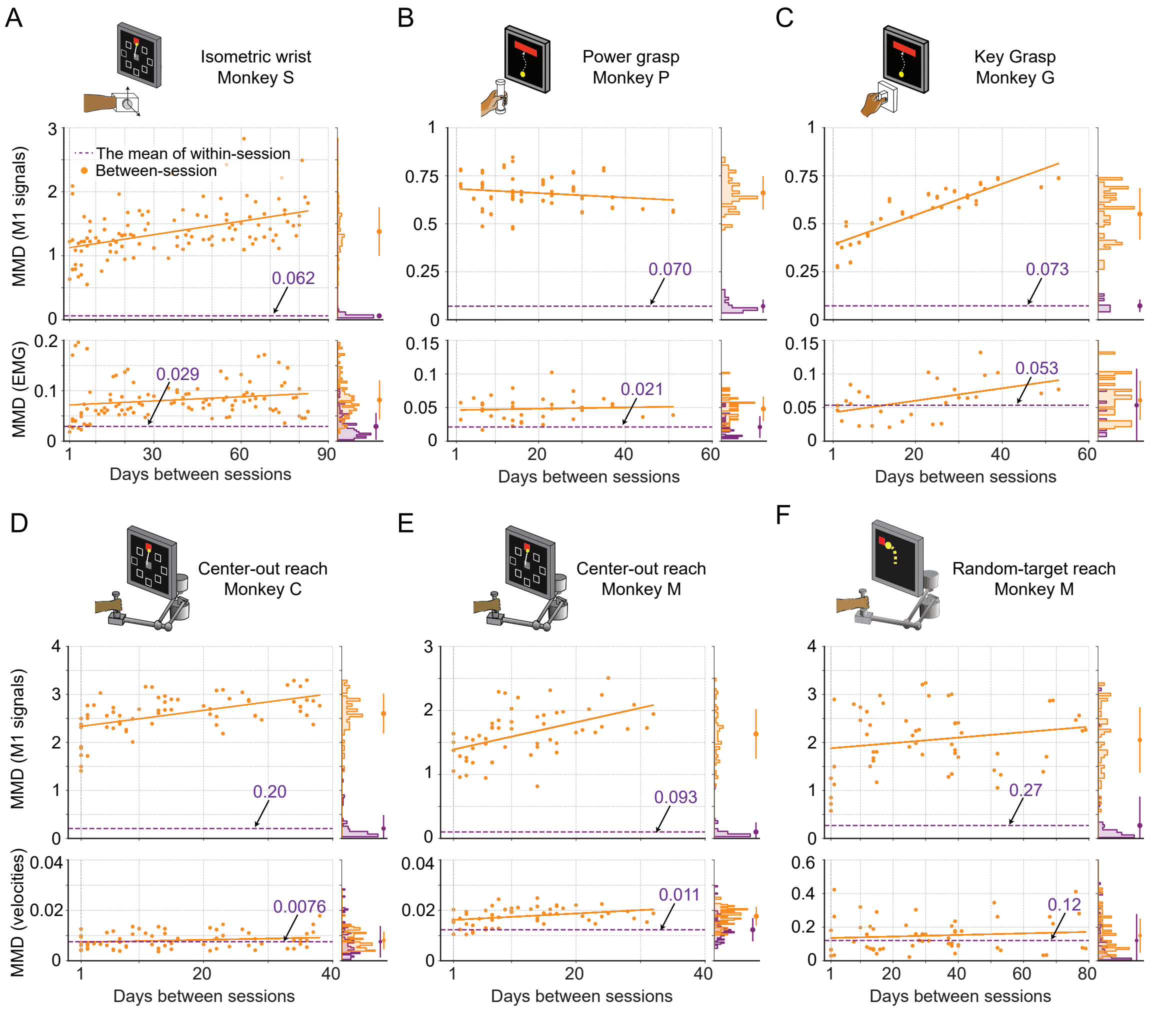


**Figure S3**. Evaluation of the stability of M1 neural signals and motor outputs over time for monkeys / tasks (besides monkey J). The stability is characterized by the discrepancy of the distributions of signals, between pairs of recording sessions in each dataset, which are measured by maximum mean discrepancy (MMD). Each subplot corresponds to a dataset: isometric wrist task of monkey S (*A*), power grasp of monkey P (*B*), key grasp of monkey G (*C*), center-out reach of monkey C (*D*) and monkey M (*E*), and random-target reach of monkey M (*F*). In each subplot, we showed the between-session MMD (orange) for M1 signals (top panel) and motor outputs (either EMG or hand velocity, bottom panel), and indicated the mean value of the within-session MMDs using a dashed purple line. The histograms for within-session and between-session MMDs were plotted on the right side of each panel, and the mean (solid dots) and standard deviation (solid lines) were shown.

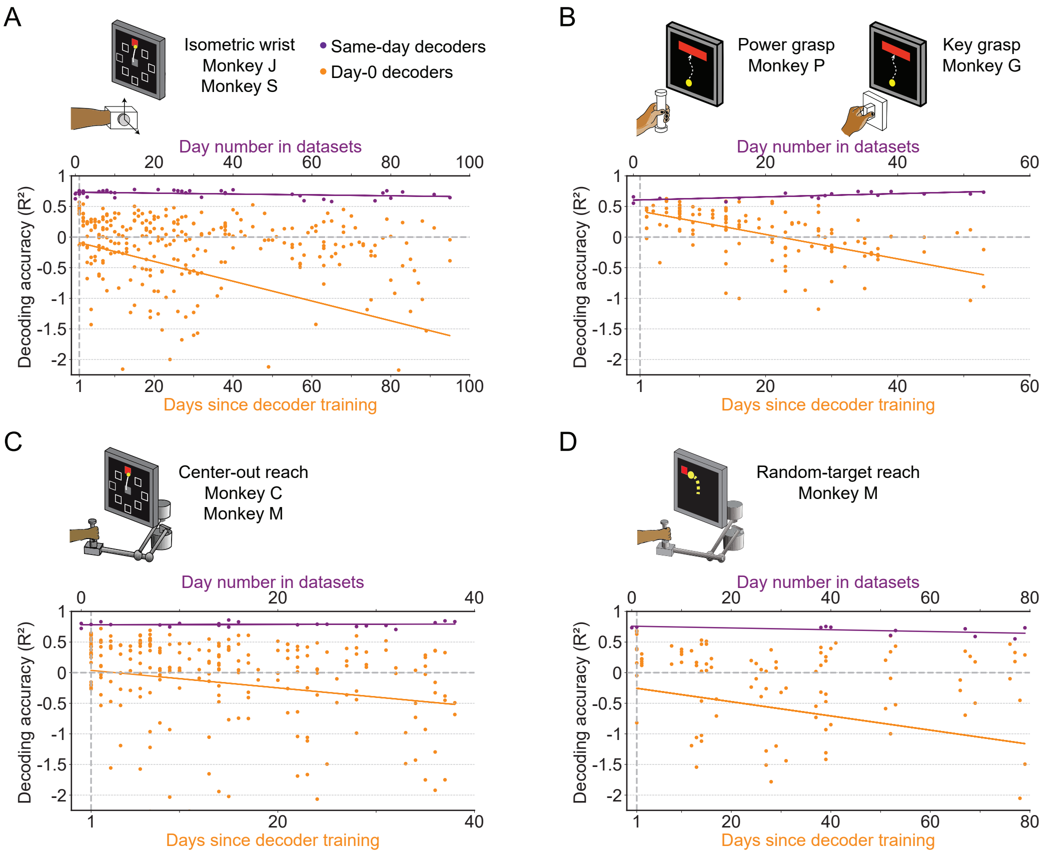


**Figure S4**. The accuracy of a well-calibrated iBCI decoder degrade over time for different behavioral tasks. We fit an iBCI decoder (Wiener filter) using the data collected on a specific day (day-0), and used this decoder to predict the motor outputs from M1 signals for all remaining days in a dataset (day-k). The performance of the decoder was evaluated by the $R^2$ value between the actual signals and the predictions. We used all available days in a dataset as the day-0 and repeated the same analysis for them. Each subplot corresponds to a behavioral task, and may contain the data from multiple monkeys: isometric wrist task of monkeys J and S (*A*), power and key grasp of monkeys P and G (*B*), center-out reach of monkeys M and C (*C*), random-target reach of monkey M (*D*). In each subplot, the $R^2$ values when using decoders to predict the motor outputs on the same day they were fit are shown (same-day decoders, purple). The x-axis on the top shows the number of the day which the recording session is on, where "0" corresponds to the earliest date in a dataset. The $R^2$ values when using decoders to predict the motor outputs on day-k are also shown (day-0 decoders, orange). The x-axis on the bottom shows the days since decoder training. The solid lines show linear fits for the $R^2$s of the same-day decoders (purple) and day-0 decoders on day-k (orange).

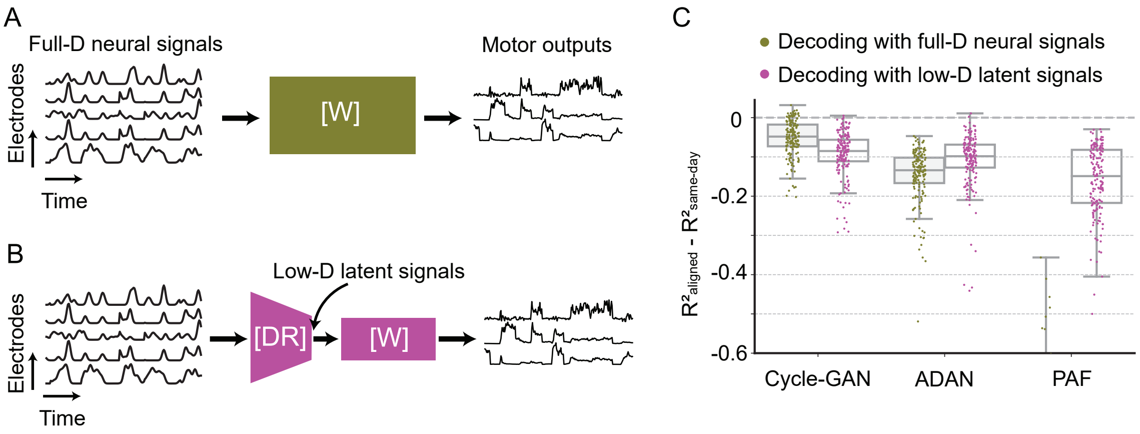


**Figure S5**. Cycle-GAN outperforms ADAN and Procrustes Alignment of Factors with both full-dimensional and low-dimensional day-0 decoder. We trained the day-0 decoders for each alignment method with either the full-dimensional firing rates (*A*) or the corresponding projections in a low-dimensional space (*B*). For the full-D decoder of ADAN and PAF, we used the reconstructed firing rates obtained from their nonlinear and linear latent space respectively. For ADAN, we used the decoder sub-network of the day-0 AE and for PAF we reversed the day-0 FA parameters to reconstruct the full-D firing rates. (*C*) Cycle-GAN outperforms ADAN and PAF with both a full-D (olive) and low-D (magenta) day-0 decoder. ADAN and PAF work better with a day-0 decoder trained on the latent signals. Note that PAF fails with a full-D decoder. For each alignment method, we computed the decoder performance drop with respect to a daily-retrained decoder (single dots: $R^2$ drop ($R^2_{aligned}$ - $R^2_{same\text{-}day}$) for days after decoder training).

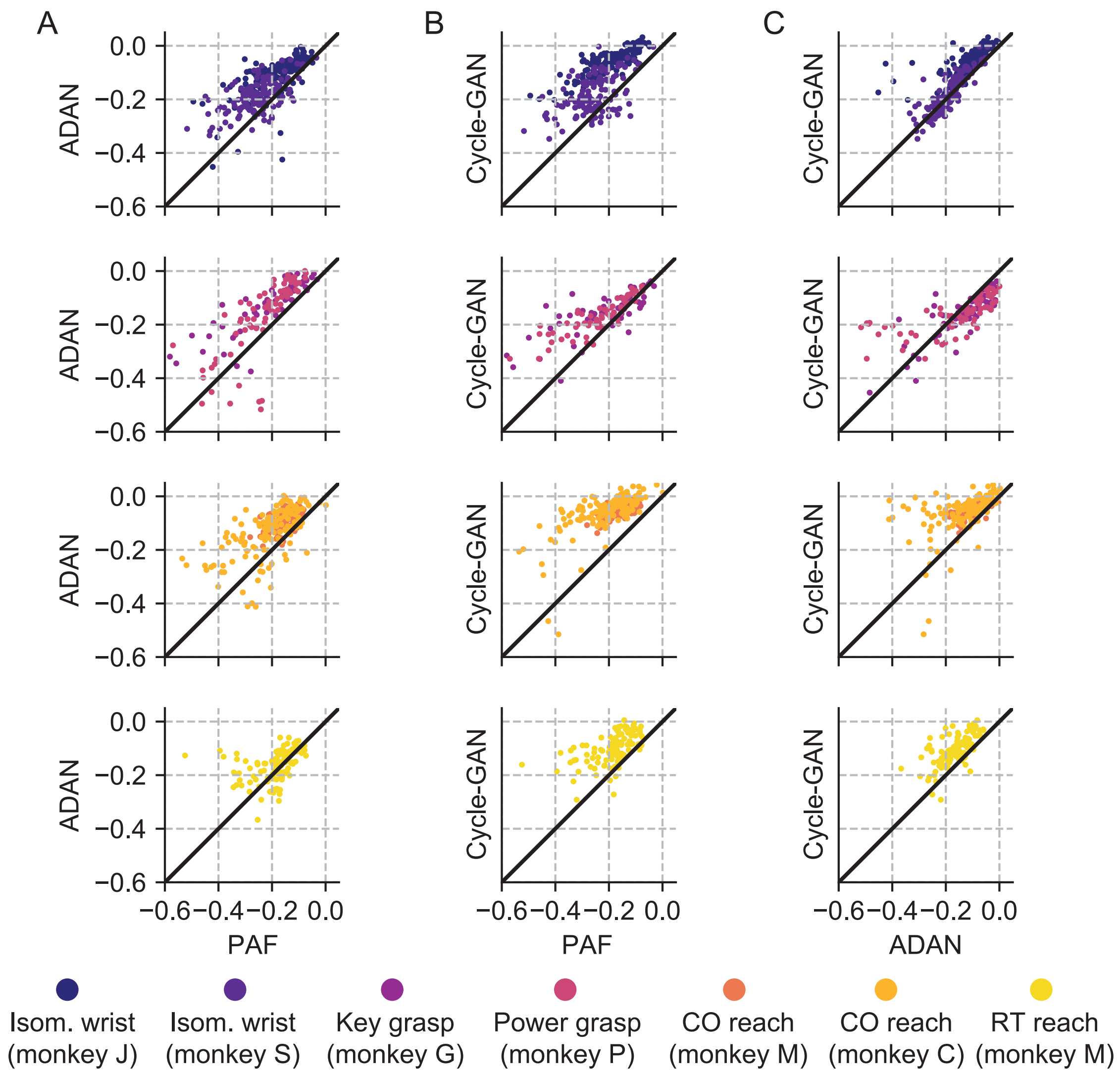


**Figure S6**. Cycle-GAN and ADAN consistently outperform Procrustes Alignment of Factors (PAF) for all experimental conditions. (*A*) ADAN vs. PAF. (*B*) Cycle-GAN vs. PAF. (*C*) Cycle-GAN vs. ADAN. Figure shows the prediction performance drop with respect to a daily recalibrated decoder ($R^2_{aligned} - R^2_{same\text{-}day}$). Each dot represents the $R^2$ drop of a given day-k. Marker colors indicate the task. Points above the unity line indicate that the aligner on the y-axis outperformed that on the x-axis. ADAN and Cycle-GAN outperform PAF for both EMG (isometric wrist, 1st row and key/power grasping, 2nd row) and kinematic (center out reaching, 3rd row and random target reaching, 4th row) decoding. Cycle-GAN performances are slightly superior to those of ADAN for the tasks where we decoded EMG (1st and 2nd row). This difference was more remarkable for the tasks where we decoded hand velocity (3rd and 4th row).

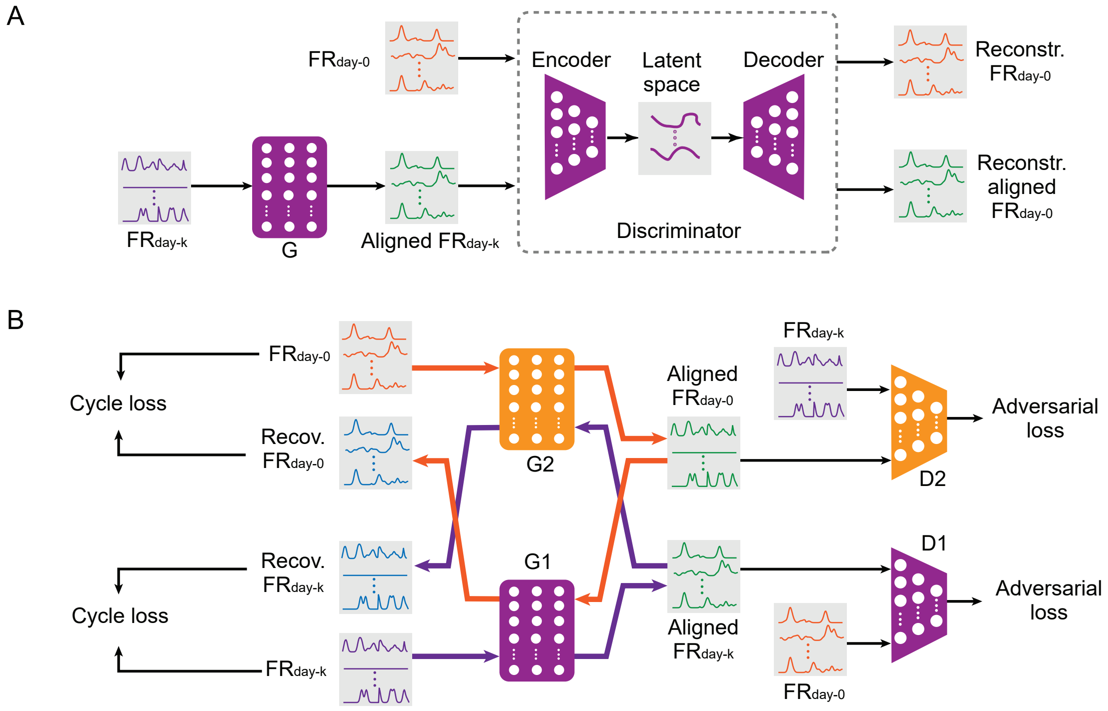


**Figure S7**. The adversarial neural networks we proposed for iBCI stabilization. (*A*) The architecture of ADAN. A feedforward neural network (G) takes the neural firing rates on day-k ($FR_{day\text{-}k}$) as the inputs, apply a transform over them to get the aligned neural firing rates (Aligned $FR_{day\text{-}k}$). Then an autoencoder takes both firing rates on day-0 ($FR_{day\text{-}0}$) and aligned $FR_{day\text{-}k}$ as the inputs, working as a discriminator. (*B*) The architecture of Cycle-GAN working as an aligner for iBCI. A feedforward neural network $G_1$ takes $FR_{day\text{-}k}$ as the inputs and gets aligned $FR_{day\text{-}k}$ after applying a transformation. Another feedforward neural network $D_1$ aims to discriminate between aligned $FR_{day\text{-}k}$ and $FR_{day\text{-}0}$, and the performance of $D_1$ contributes to the "adversarial loss". Another pair of feedforward neural networks $G_2$ and $D_2$ functions in the same way as $G_1$ and $D_1$, but aims to convert $FR_{day\text{-}0}$ into a form like $FR_{day\text{-}k}$. The outputs of $G_1$, the aligned $FR_{day\text{-}k}$, are also sent to $G_2$, which aims to recover the original $FR_{day\text{-}k}$ from them. The discrepancy between the real $FR_{day\text{-}k}$ and the recovered $FR_{day\text{-}k}$ contributes to the "cycle loss". The purple arrows highlight this workflow, forming a "cycle" like structure. Likewise, the orange arrows highlight the workflow of another cycle.

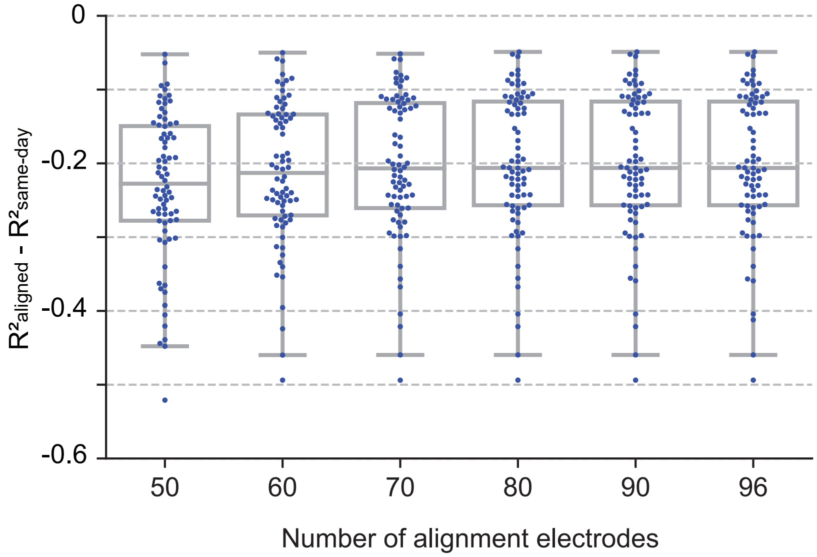


**Figure S8**. PAF performs better when a greater number of electrodes used for alignment is selected. The effect of number of electrodes for alignment on performance is shown. For each number to be tested, we computed the decoder performance drop with respect to a daily-retrained decoder (single dots: $R^2$ drop ($R^2_{aligned}$ - $R^2_{same\text{-}day}$) for days after decoder training). The performance of the day-0 decoders after alignment increases with the number of electrodes used for alignment and reaches a plateau at 70.
